# Negative regulation of the Rcs phosphorelay via IgaA contact with the RcsD phosphotransfer protein

**DOI:** 10.1101/2020.01.14.906271

**Authors:** Erin A. Wall, Nadim Majdalani, Susan Gottesman

## Abstract

Two-component systems and phosphorelays play central roles in the ability of bacteria to rapidly respond to changing environments. In *E. coli* and related enterobacteria, the complex Rcs phosphorelay is a critical player in changing bacterial behavior in response to antimicrobial peptides, beta-lactam antibiotics, and other challenges to the cell surface. The Rcs system is unusual in that IgaA, an inner membrane protein, is essential due to its negative regulation of the RcsC/RcsD/RcsB phosphorelay. While it has previously been shown that IgaA transduces signals from the outer membrane lipoprotein RcsF, how it interacts with the phosphorelay was unknown. Here we use in vivo interaction assays and genetic dissection of the critical proteins to demonstrate that IgaA interacts with the phosphorelay protein RcsD, and that this interaction is necessary for regulation. Interactions in periplasmic domains of these two proteins anchor repression of signaling. However, the signaling response depends on a weaker interaction between cytoplasmic loop 1 of IgaA and a truncated PAS domain in RcsD. A point mutation in the PAS domain increases interactions between the two proteins and is sufficient to abolish induction of this phosphorelay. RcsC, the histidine kinase that initiates phosphotransfer through the phosphorelay, appears to be indirectly regulated by IgaA via the contacts with RcsD. Unlike RcsD, and unlike many other histidine kinases, the periplasmic domain of RcsC is not necessary for the response to inducing signals. The multiple contacts between IgaA and RcsD form a poised sensing system, preventing over-activation of this apparently toxic phosphorelay but allowing it to be rapidly and quantitatively responsive to signals.

**Author Summary:** The Rcs phosphorelay plays a central role in allowing enterobacteria to sense and respond to antibiotics, host-produced antimicrobials, and interactions with surfaces. A unique negative regulator, IgaA, keeps signaling from this pathway under control when it is not needed, but how it controls the phosphorelay has been unclear. We define a set of critical interactions between IgaA and the phosphotransfer protein RcsD. A periplasmic contact between IgaA and RcsD provides a necessary inhibition of Rcs signaling, modulated further by regulated interactions in the cytoplasmic domains of each protein. This multipartite interaction provides a sensitive regulatory switch.

## Introduction

Bacteria must constantly monitor their cell wall and envelope integrity to withstand environmental insult. Osmotic stress, redox stress and envelope disruption demand that the bacterium remodel its exterior to provide protection, often in the form of capsular polysaccharide. Enterobacterales use the Rcs phosphorelay to integrate complex signals from the outer membrane and periplasm, changing gene regulation in response to stress [1, 2]. The Rcs phosphorelay is a complex signal transduction pathway, involving an outer membrane lipoprotein (RcsF) and three inner membrane proteins (IgaA, RcsC and RcsD), leading to changes in the phosphorylation of the transcriptional regulator (RcsB). The Rcs phosphorelay regulates production of virulence-associated capsules as well as motility and the expression of many stress-related genes.

Signaling through the pathway is complex, and not fully understood. Briefly, outer membrane stress such as cationic polypeptides or cell wall stresses such as beta-lactams cause RcsF to change its interaction with IgaA (originally identified in *Salmonella* and named for intracellular growth attenuation; the *E. coli* version of this gene, *yrfF*, is referred to here as IgaA). The activated RcsF/IgaA interaction allows the hybrid histidine kinase RcsC to auto-phosphorylate and then pass phosphate to phosphorelay protein RcsD, a process studied here, which passes it to response regulator RcsB (Figure 1A). Over-signaling through the phosphorelay leads to cell death, possibly because of the global nature of the RcsB regulon. IgaA is essential because of its role as a gating/braking mechanism for the phosphorelay. Deletion of IgaA is only possible in cells containing mutations in RcsC, RcsD, or RcsB. For this reason, the poorly understood IgaA mechanism of action is of key interest. Multiple studies have focused on the interaction of RcsF with IgaA when cell wall stress is detected [3–7], but the downstream action of IgaA is less well understood [8]. In this work we define RcsD as the direct binding partner of IgaA and define the regions in RcsD that are critical for interaction with IgaA. Production of RcsD variants that are deficient in IgaA binding cause massive over-signaling, mucoidy, and often cell death, consistent with phenotypes seen upon loss of IgaA itself. These results contrast with previous assumptions that IgaA was likely to directly interact with and regulate the histidine kinase RcsC, because RcsC initiates the phosphorelay.

**Figure 1:**
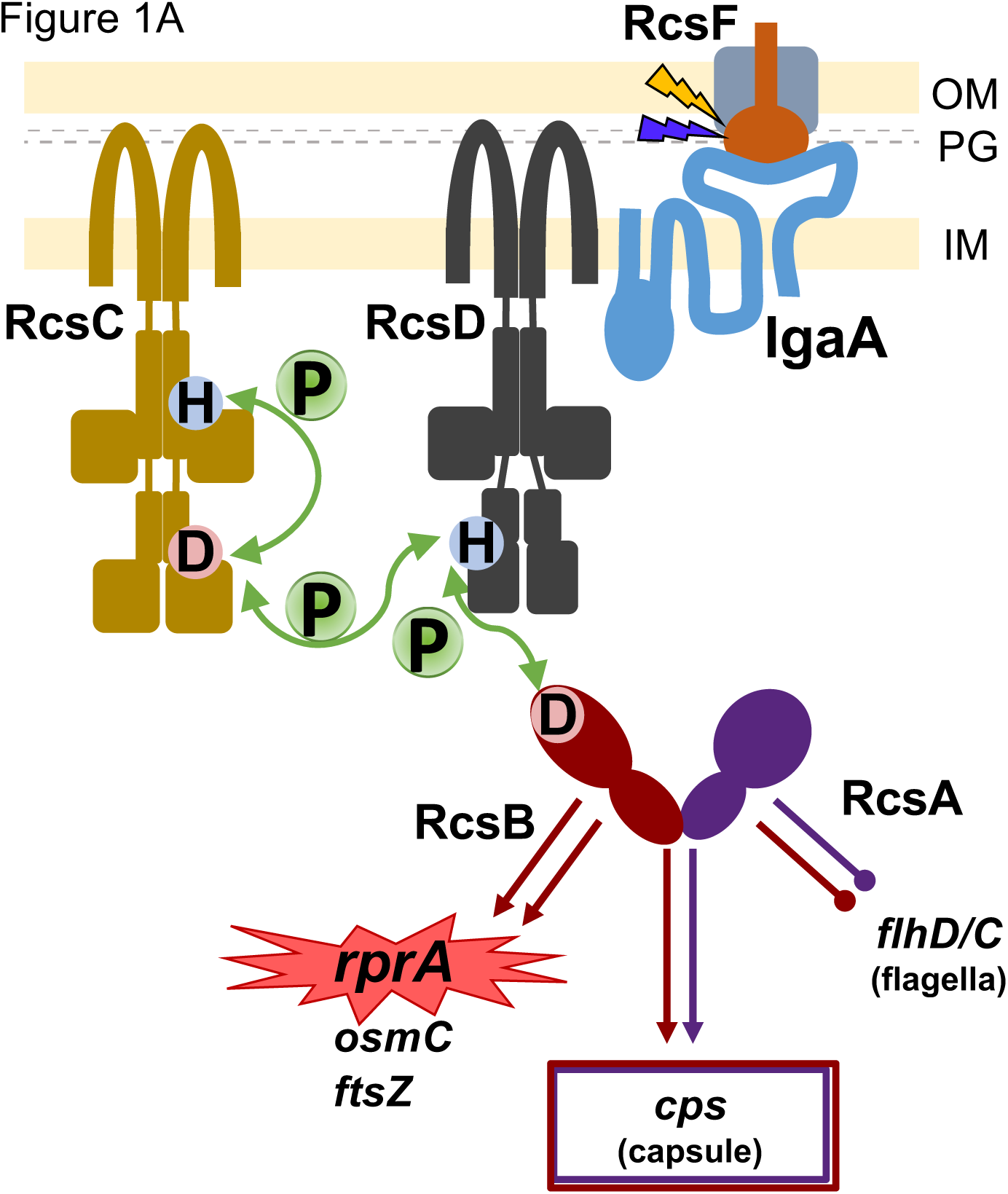
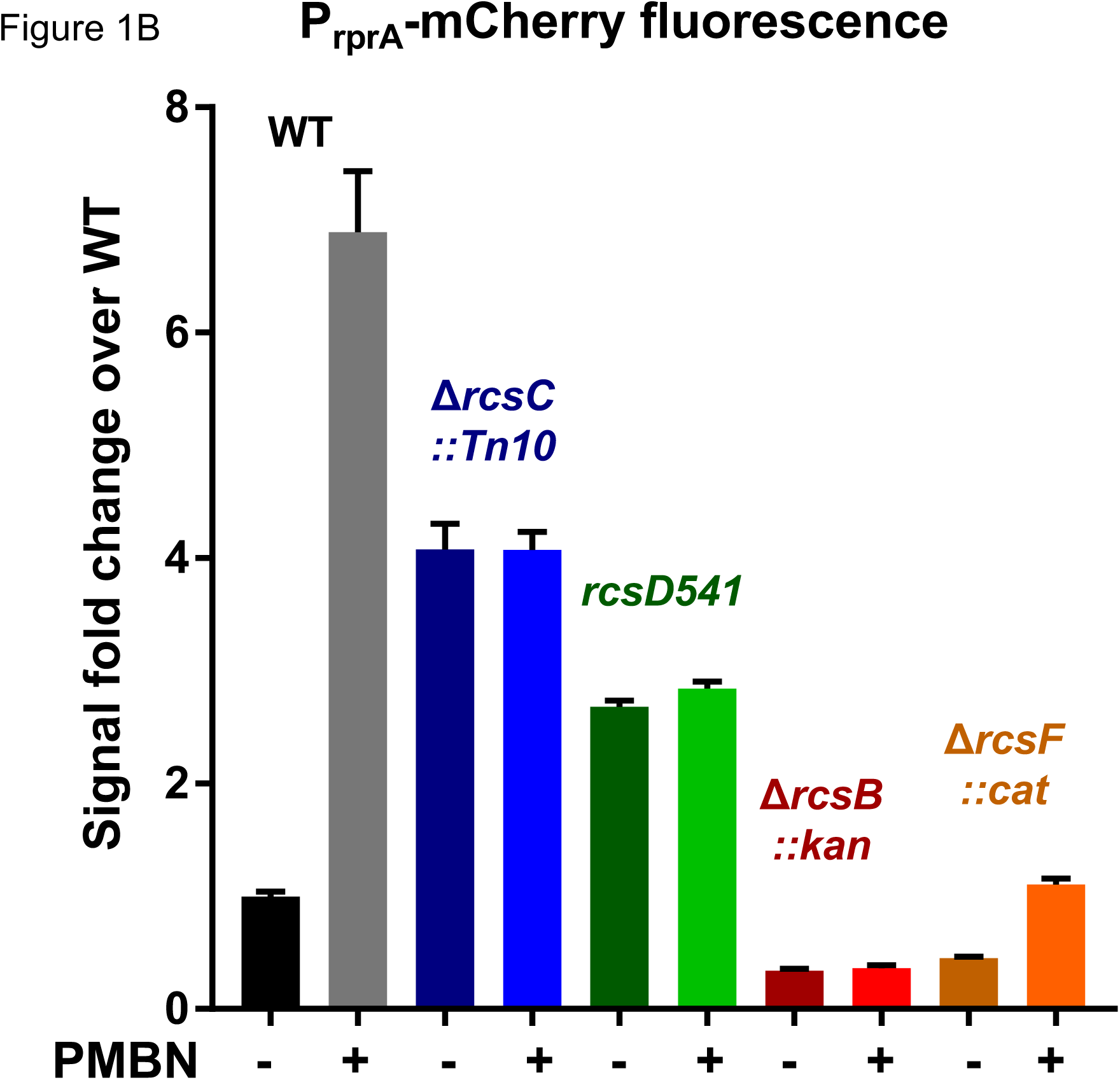
Signaling via the Rcs Phosphorelay. A. The six proteins of the Rcs Phosphorelay are shown schematically (not to scale; described in detail in [1]). RcsF is positioned in the outer membrane, associated with outer membrane porins (OMPs). Most described treatments that induce the phosphorelay require RcsF for activation and thus it is shown as a key sensor for both outer membrane stress (represented by a gold lightning bolt) and periplasmic or peptidoglycan stress (blue lightning bolt). IgaA is a five-pass inner membrane protein that serves as a brake on the phosphorelay; it communicates with RcsF across the periplasm. Current models suggest that, upon stress signaling, RcsF contacts or changes contacts with IgaA, leading to de-repression of the phosphorelay. RcsC is induced to autophosphorylate and pass phosphate from its active site His 479 to its REC domain Asp 875. The phosphate is then passed to His 842 on the RcsD histidine phosphotransfer domain, which then passes to the RcsB REC domain Asp 56. Phosphorylated RcsB both homodimerizes and heterodimerizes with RcsA to regulate many genes, notably repressing flagellar synthesis, inducing capsule synthesis, and inducing the sRNA RprA. The red highlight around *rprA* indicates that an *rprA* promoter fusion to mCherry (P_rprA_-mCherry) is used throughout this work to evaluate activation of the phosphorelay. Note that as with many phosphorelays of this family, phosphate can also flow in reverse from RcsB towards RcsC. IgaA is shown closest to RcsD, as discussed in this paper. B. The promoter of the sRNA RprA was fused to mCherry to create a reporter for Rcs activation (P_rprA_-mCherry), demonstrating sensitivity and a wide dynamic range. Activity of wild type cells (black, EAW8) was compared to Δ*rcsC::Tn10* (blue, EAW18), *rcsD541* (green, EAW19), Δ*rcsB::kan* (red, EAW31) *and* Δ*rcsF::chl* (orange, EAW32). All strains were also tested with polymyxin B nonapeptide (PMBN) at 20ug/mL. Cells were grown in MOPs minimal glucose for the P_rprA_-mCherry assay; signal at OD_600_ 0.4 is shown. Details of the assay and cell growth are shown in S1A-C Fig and described in Materials and Methods.

## Results

### A sensitive and flexible assay for the Rcs phosphorelay

We have revisited the Rcs signaling pathway using a newly developed sensitive in vivo fluorescent reporter assay. RprA is a small RNA that is a sensitive, specific target of Rcs regulation. RprA levels are nearly undetectable in the absence of RcsB, its direct transcriptional activator, and increase in cells in which the Rcs system is activated, for instance by constitutively active *rcsC* mutations [9]. A *rprA* transcriptional fusion to mCherry allows continuous detection of a wide range of Rcs activation levels, using a plate reader. This growth and fluorescence assay can be viewed as a function of fluorescence over average OD, showing a large, early change in slope when a given strain is induced to respond to an Rcs stimulus (Fig S1A-C). Bar graphs of fluorescence at OD_600_ 0.4. Figure 1B shows the increase in signal when wild-type cells are exposed to a small molecule stimulator of Rcs signaling, the non-toxic cationic polymyxin B nonapeptide (PMBN). PMBN stimulus is a useful indicator of pathway status; normal signaling yields a measured level of response to PMBN, while pathway disruptions (modification or deletion of pathway components) cause dampened or loss of PMBN response. Cells deleted for *rcsB* lose all signal, including the low basal level seen in the absence of PMBN (Fig 1B, S1A-C Fig). The absence of RcsF lowers overall signal (compare Δ*rcsF*::chl, - PMBN, to WT, - PMBN); this decrease has been reported before [3, 10–12], but is particularly clear with this assay. Lack of RcsF also greatly dampens response to an outer membrane stress like PMBN (Fig 1B). It is known that Rcs signaling can be induced in the absence of RcsF, so the small activation of the *rcsF*::chl strain in the presence of PMBN is possibly stimulating Rcs in this (still unexplored) manner [10, 13]; (Majdalani et al, unpublished).

The hybrid histidine kinase RcsC and the phosphorelay protein RcsD play both positive and negative roles in regulation of RcsB activity. Loss of RcsC or RcsD blocks the response to PMBN, but also lead to significantly higher levels of P_rprA_-mCherry in the absence of normal inducing signal (Fig 1B). This is consistent with previous work, in which expression of an P*_rprA_- lac* reporter was increased upon deletion of *rcsC* or *rcsD* [9, 11]. Deletion of *rcsC* is thought to cause the loss of ability to de-phosphorylate RcsB that has acquired phosphate from other sources [14–16]. In our assay conditions, the *rcsC* deletion strain produces a signal that is 3-4 fold above WT.

The *rcsD* deletion shown, *rcsD*541, increased P_rprA_-mCherry expression in a manner similar to the *rcsC* deletion. *rcsD* is encoded upstream of *rcsB*, with the major promoters for *rcsB* inside the *rcsD* coding region [17]. This affects the way *rcsD* deletion alleles can be constructed. In addition, in both *Salmonella* and *E. coli*, some transcripts from the *rcsD* promoter may continue through to *rcsB*, though apparently at a much lower level [17, 18]. Four different *rcsD* alleles were examined, each designed to leave the *rcsB* promoters intact (S1D Fig). These include *rcsD* carrying an H842A mutation in the phosphotransfer domain active site, as well as *rcsD* containing stop codons after codon 841 (*rcsD*841*). The overall amount of P_rprA_-mCherry expression seems to differ modestly between *rcsD541* and *rcsD543*, even though the deleted regions in both alleles share the same boundaries. *rcsD*541 also gave higher P_rprA_-mCherry signal than the point mutants (S1D Fig). When checked by western blot with a polyclonal RcsD antibody, it is evident that *rcsD*541, *rcsD*543 and *rcsD*841* are all devoid of detectable RcsD (S1E Fig). As previously seen [11], *rcsD*541 and *rcsD*543 had no significant effect on RcsB levels, nor did *rcsD* H842A and *rcsD*841* (S1E Fig, right panel).

The somewhat increased level of P_rprA_-mCherry common to all *rcsD* and *rcsC* strains is likely in part due to phosphorylation of RcsB by the small molecule acetyl phosphate (AcP), in the absence of the dephosphorylation carried out by RcsD and RcsC [15, 19]. The influence of AcP can be demonstrated in an *ackA* deletion strain that builds up large amounts of AcP [15] (S1F Fig). While WT cells showed a modest increase in signal in the *ackA* mutant (compare black and gray bars), all tested *rcsC* and *rcsD* mutations produced high levels of P_rprA_-mCherry I the *ackA* mutant, consistent with failure to dephosphorylate RcsB (S1F Fig). The increase in signal is fully dependent upon RcsB (last bar in graph, S1F).

### IgaA and RcsD interact directly

We began interrogating how IgaA might interfere with Rcs signaling by examining the interactions of IgaA with downstream members of the phosphorelay, using the bacterial adenylate cyclase two hybrid assay (BACTH). In this assay, cells produce beta-galactosidase when Bordetella adenylate cyclase fragments (T18 and T25, Cya) are reconstituted by fusion to interacting proteins in a cyclase-defective host [20, 21]. IgaA interacted robustly with RcsD in two orientations (IgaA-T18/RcsD-T25 and IgaA-T25/RcsD-T18), expressing beta-galactosidase activity approximately 20-fold greater than either fusion paired with an empty cognate vector, the standard background control (Fig 2A; S2A Fig). This interaction occurred irrespective of the chromosomal presence or absence of other Rcs members (S2B,C Fig). However, no significant interaction was detected between IgaA and RcsC in parallel assays (Fig 2A, S2B,D Fig).

**Figure 2:**
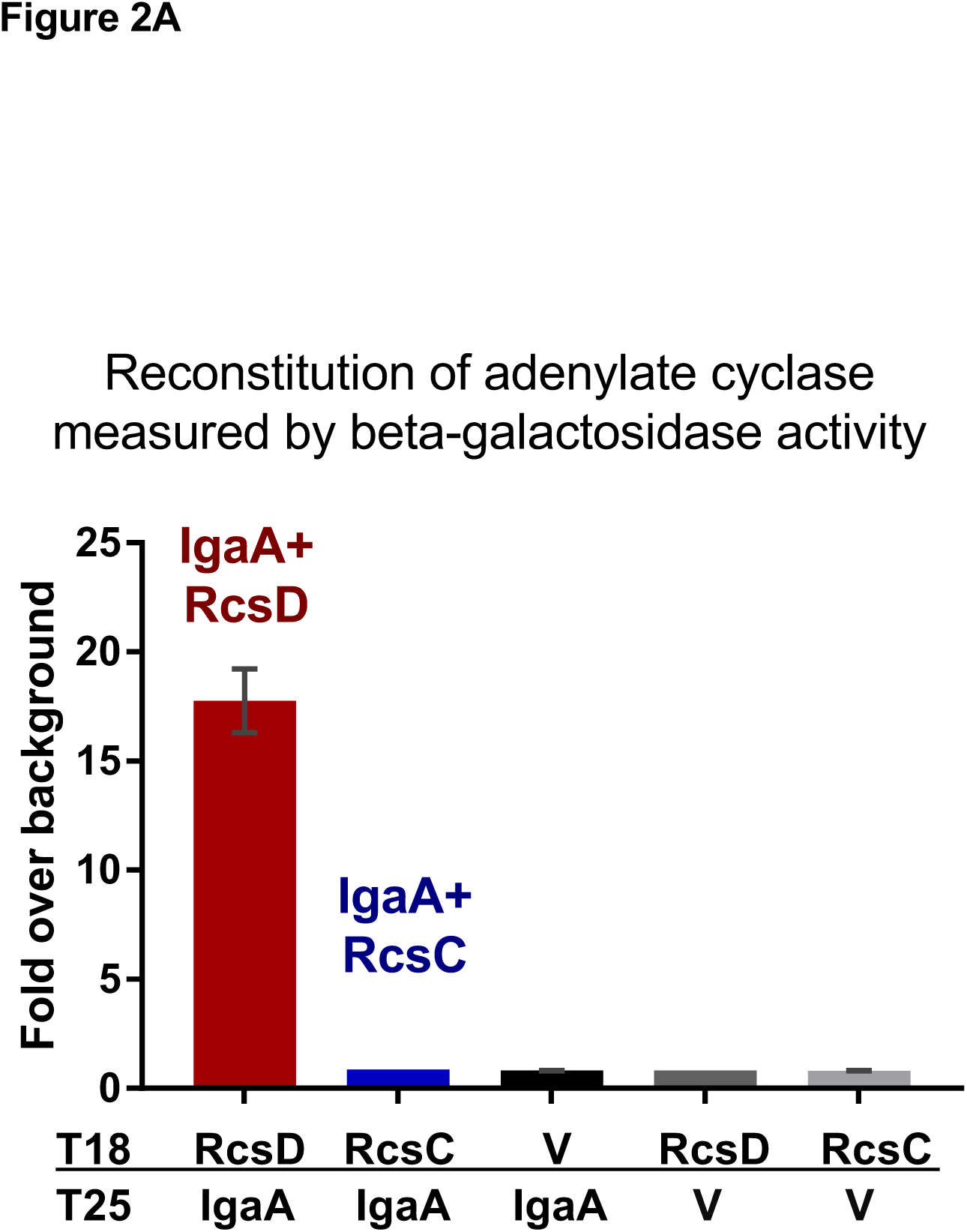
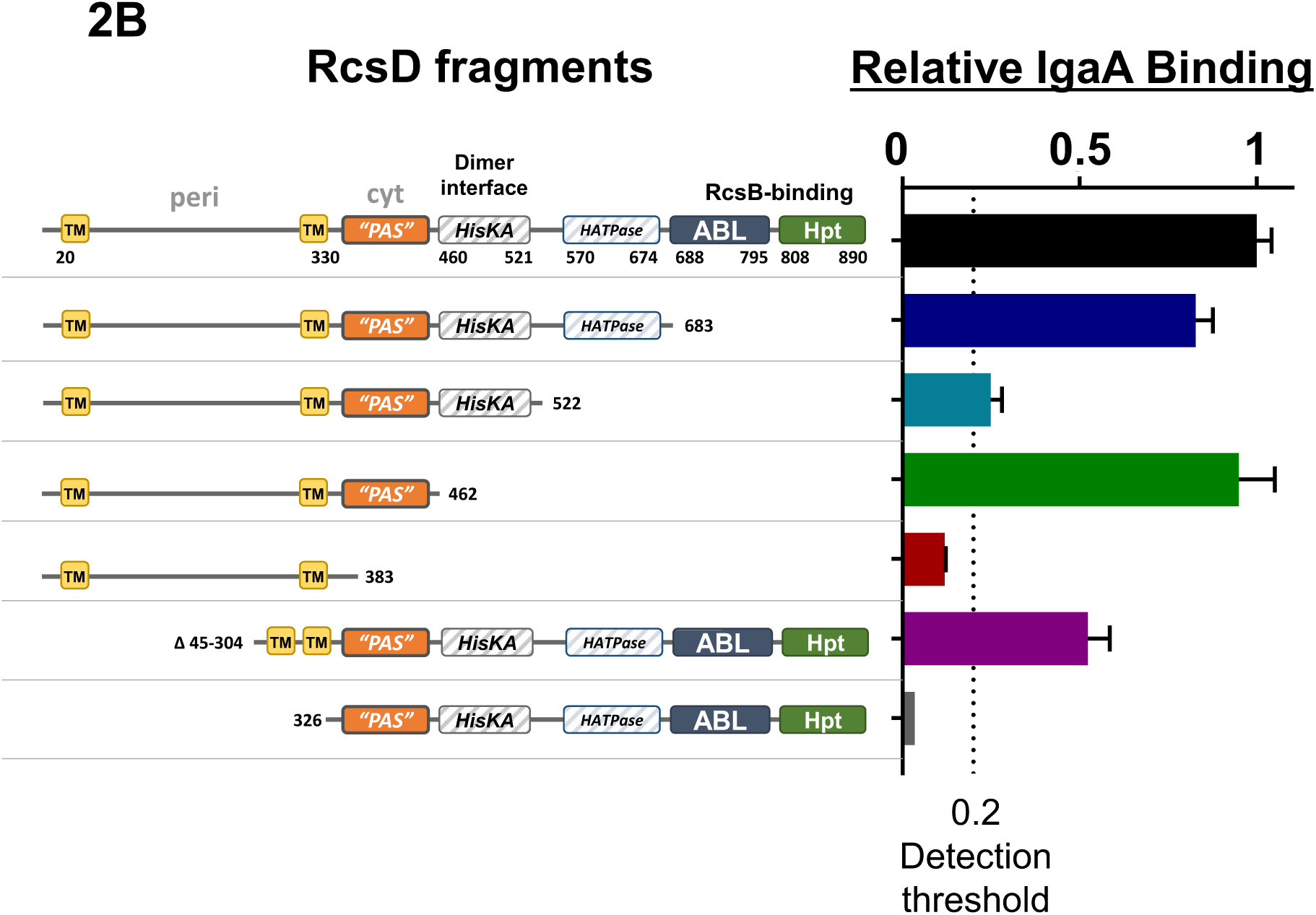
Interaction of IgaA with RcsD. A. Relative beta galactosidase activity was measured in *cyaA* deficient cells (strain BTH101) containing a dual plasmid system encoding the T18 and T25 domains of adenylate cyclase. Each protein fusion plasmid paired with its cognate vector produces very little activity (gray bars). Error bars (some too small to be visible) represent standard deviation of three assays. Fusions present are IgaA-T25, RcsD-T18, and RcsC-T18. The opposite orientation, tests of roles of other Rcs members on the interaction, and Miller units are shown in S2A-D Fig. B. Relative ratio of RcsD fragment binding to IgaA was determined by comparing the interaction of RcsD truncations to the interaction between full length RcsD and IgaA, set to 1. In most cases the IgaA/RcsD interaction is 20x over background, usually 1000 Miller units to 50 Miller units for the background control. The dotted line at y=0.2 represents the threshold for interaction detection, 4x over background signal. All measurements were taken in BTH101. Plasmids present were pEAW1 (IgaA-T18), pEAW8 (RcsD-T25), pEAW8b (RcsD_683_-T25), pEAW8α (RcsD_522_-T25), pEAW8m2 (RcsD_462_-T25), pEAW8m (RcsD_383_-T25), pEAW8peri (RcsD_Δ45-304_-T25), pEAW8s (RcsD_326-C_-T25).

The IgaA, RcsD, and RcsC fused to Cya fragments were all tested for their ability to function in the Rcs phosphorelay and were found to be functional (see Materials and Methods, S2E Fig); blots for the proteins showed the expected bands (S2F Fig). Therefore, lack of interaction by RcsC in the bacterial two-hybrid assay cannot be attributed to significant misfolding or lack of protein. These results suggest that IgaA interacts with RcsD but does not interact with RcsC in this assay.

Regions in RcsD necessary and sufficient for interaction with IgaA were defined in the bacterial two-hybrid assay (Fig 2B). Using C-terminally truncated RcsD constructs, we found that IgaA bound just as well to RcsD_N-683_ (RcsD without the ABL and Hpt domains) as it did to wild type. It also bound to RcsD_N-522_ (no HATPase, ABL or Hpt domains), but less strongly than to wild type or RcsD_N-683_. The interactions were unaffected by the presence of RcsD in the chromosome (S2G Fig). Strikingly, RcsD_N-462_ also bound IgaA almost as well as full length RcsD (Fig. 2B). This region is predicted to contain an incomplete Per-Arndt-Sim (PAS) domain (shown on the graphic as “PAS” to emphasize that it is not a complete PAS domain), which has been associated with signal detection in sensor histidine kinases [22]. A further truncation, removing most of the cytoplasmic regions upstream of the HATPase domain (RcsD_N-383_), did not produce a measurable IgaA interaction, suggesting a critical role for at least some of the cytoplasmic domain. A fully cytoplasmic RcsD construct (RcsD_326-C_) that began directly after the membrane-bound portion and included the rest of the RcsD C-terminus also failed to show any detectable interaction with IgaA. Finally, a construct in which the periplasmic region was deleted (RcsD_Δ45-304_) still interacts with IgaA, although only at a level of about half that seen with the WT construct (Fig 2B).

These results are most consistent with RcsD interactions with IgaA both within the trans-membrane/periplasmic portion of RcsD and within the initial cytoplasmic regions (bounded perhaps by residue 462), with neither interaction sufficient for a full signal in this two-hybrid assay. Intriguingly, the constructs that gave positive interactions (RcsD_N-522_, RcsD_N-683_ and RcsD_N-462_, but not RcsD_N-383_ and RcsD_326-C_) also caused mucoidy, a reflection of activation of the Rcs phosphorelay, in the cloning strain (Stellar *E. coli*, Clontech), which is wild-type for all genes of the Rcs phosphorelay. This was further examined, using the P_rprA_-mCherry assay.

### Titration of IgaA by overexpression of truncated RcsD

If IgaA repression of Rcs signaling depends on the direct interaction of RcsD with IgaA, suggested by the BACTH results, overproduction of the regions of RcsD sufficient for this interaction may titrate IgaA away from the chromosomally-encoded RcsD, allowing unregulated signaling through the Rcs phosphorelay. The same RcsD fragments studied in the BACTH experiments were cloned without tags into a pBAD24 plasmid, under the control of the arabinose-inducible pBAD promoter. In this plasmid in the absence of arabinose, RcsD is expressed levels sufficient to complement the modest increase in signal seen in an *rcsD* mutant (Fig 3A, right panel; compare RcsD to V); the protein is significantly overexpressed in the presence of arabinose. These plasmids were assayed in both *rcsD*^+^ and *rcsD*541 strains containing the P_rprA_-mCherry reporter fusion. We would expect the WT strain to be active for the reporter when the Rcs phosphorelay is activated. Indeed, in the *rcsD*^+^ host, overproduction after arabinose induction of RcsD fragments capable of interacting with IgaA (RcsD_N-683_, RcsD_N-525_ and RcsD_N-462_) activated the P_rprA_-mCherry fusion, while RcsD_N-383_, which was negative for interaction with IgaA (Fig 2B), did not (Fig 3A, S3A Fig). Significant cellular growth arrest and lysis occurred when RcsD_N-683_ or RcsD_N-525_ were overproduced (in the presence of arabinose), making them difficult to compare quantitatively to non-lysed cells. In S3A Fig, their activation at lower ODs (before lysis) is shown, and fluorescence graphed as a function of the OD throughout growth is shown.

**Figure 3:**
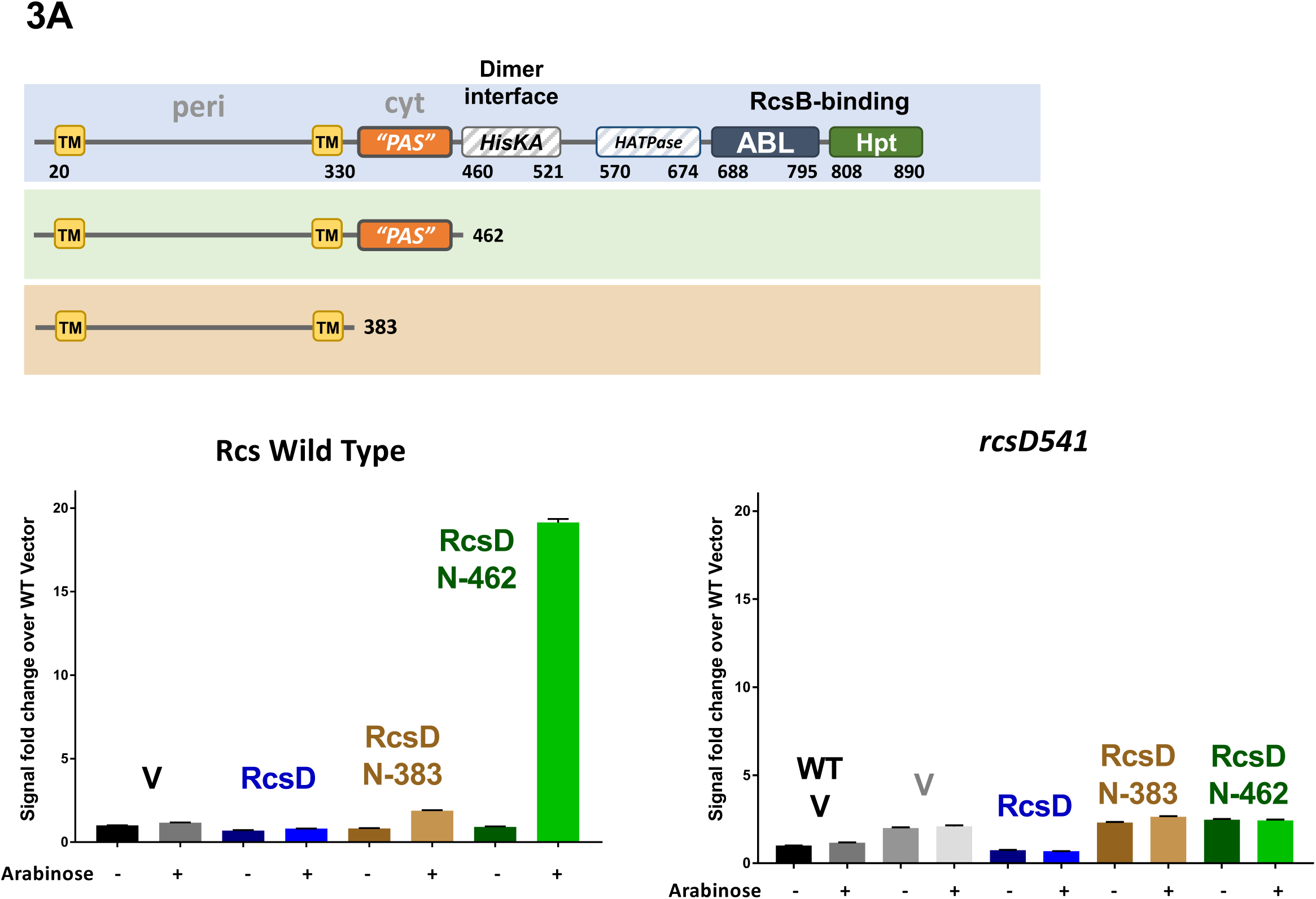
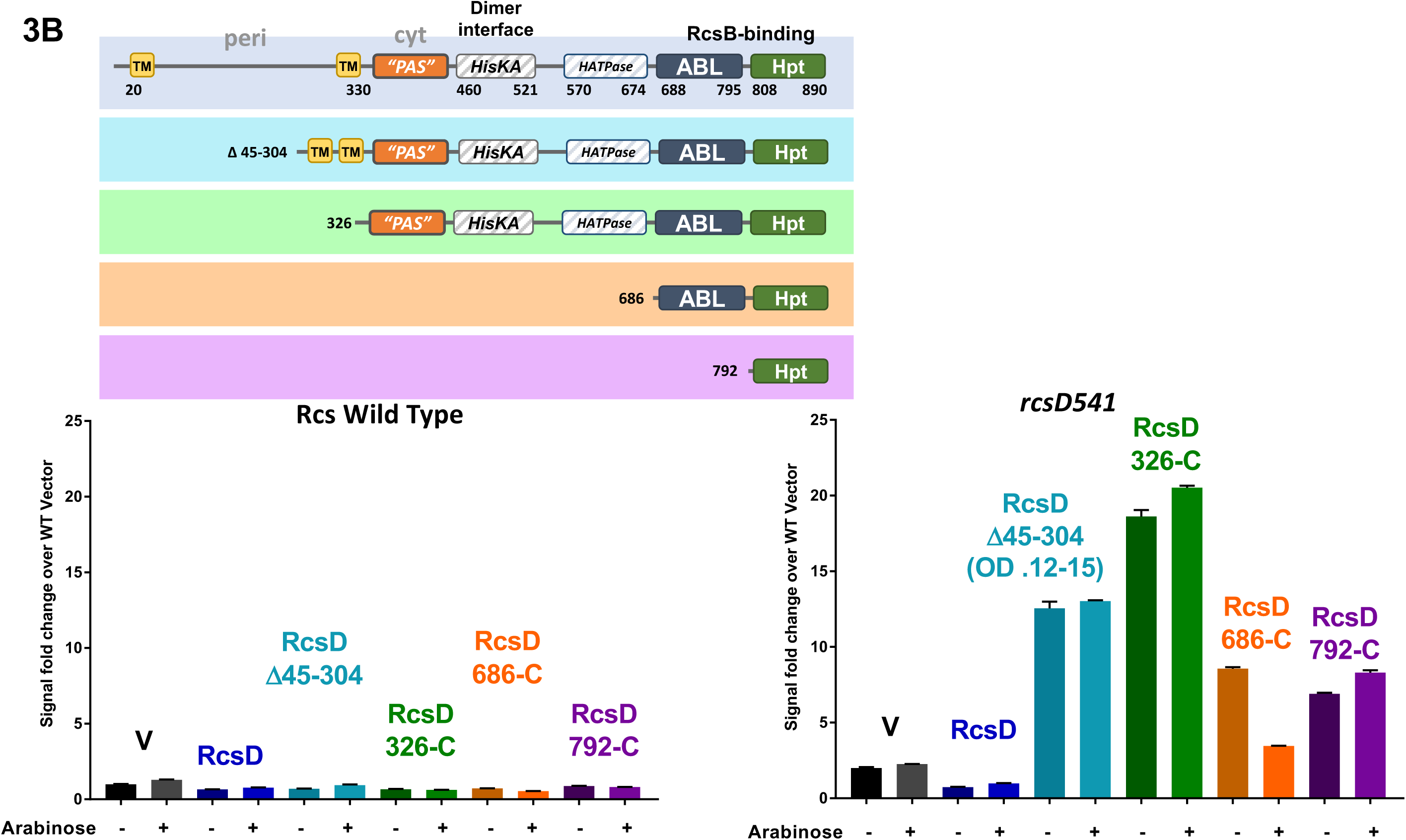
Activity of truncated RcsD proteins. A. WT (EAW8) and *rcsD541* (EAW19) mutant cells carrying the P_rprA_-mCherry fusion were transformed with pBAD24-derived plasmids encoding RcsD or truncated pieces of RcsD, grown in MOPS glucose with ampicillin (-arabinose) or MOPS glycerol with ampicillin, with 0.02% arabinose (+ arabinose). Fluorescence values for each strain are shown at OD_600_ 0.4. RcsD truncations used are shown, with color-coding: black: V (vector, pBAD24); blue: RcsD (pEAW11); brown: RcsD_N-383_ (pEAW11m); green: RcsD_N-462_ (pEAW11m2). Fluorescence as a function of OD_600_ and additional related plasmids in the same strains are shown in S3A Fig; results in other strain backgrounds are shown in S3B-C Fig. B. Experiments as in panel A, in *rcsD*^+^ (WT; EAW8) and *rcsD*541 (EAW19) hosts. The constructs are color-coded as follows: black: vector (pBAD24), blue: full length RcsD (pEAW11), cyan: RcsD_Δ45-304_ (pEAW11peri), green: RcsD_326-C_ (pEAW11s), orange: RcsD_686-C_ (pEAW11c), purple: RcsD_792-C_ (pEAW11d). Note that for the *rcsD*541 cells carrying RcsD_Δ45-304_, the value shown is at a low OD, the total achieved within 6 hours.

The ability of fragments to activate in an *rcsD*^+^ host correlates well with ability to interact with IgaA in the bacterial two-hybrid assay, consistent with the model that activation by overproduction is a result of binding to and titrating IgaA, freeing the wild-type RcsD to signal. These results reinforce the conclusion from the BACTH assays that at least two regions of RcsD interact with IgaA. Both regions are necessary for efficient titration, one region presumably between 1-383 (encompasses trans-membrane and periplasmic region), present in all the titrating plasmids, but not sufficient for titration, and a second region between 383 and 462 (including the incomplete PAS-like domain).

Note that overexpressing full-length RcsD did not induce mucoidy or signaling. Since the full-length protein has all regions that should bind and titrate IgaA, this result suggests the possibility that the C-terminal domains of RcsD, missing in all the activating/titrating truncations, might play a negative role that blocks or is epistatic to the titration seen with the truncated plasmids. This is consistent with the report that the ABL domain can bind and inhibit phosphorylation of RcsB [23], and is confirmed below.

The RcsD Hpt domain is necessary for transmitting a signal from the RcsC response regulator domain to RcsB [24]. Therefore, we expected that plasmids expressing truncated RcsD constructs that lack the Hpt domain would be completely unable to activate the phosphorelay in a strain mutant for *rcsD*. In the *rcsD541* mutant allele background, the basal level of expression is above that in a WT host (Fig 1B, Fig 3A, compare V to WT V); a plasmid expressing the intact RcsD reduces P_rprA_-mCherry expression to levels comparable to the WT strain, consistent with complementation of the *rcsD*541 mutant (Fig. 3A). The activating fragment RcsD_N-462_ did not induce significant P_rprA_-mCherry activity in this host, consistent with expectation, since it does not contain the Hpt domain (Fig 3A).

Unexpectedly, cells expressing somewhat longer RcsD fragments, truncations RcsD_N-522_ and RcsD_N-683_, both missing the Hpt domain, were able to significantly increase signal when induced in the *rcsD*541 host (S3A Fig). However, these same plasmids caused lysis in WT and *rcsD*541 cells (S3A Fig). These constructs contain regions of RcsD that are not well understood, including an ancestral histidine kinase structure between residues 462 and 683. To further investigate the basis for this unexpected signaling, the same plasmids were tested in three additional *rcsD* mutants (S3B Fig). In a strain with a chromosomal mutation in the phosphotransfer active site, RcsD H842A,or in a strain containing a stop codon at residue 842, right before the active site (*rcsD*841*), the RcsD_N-522_ and RcsD_N-683_ truncations did not raise the level of P_rprA_-mCherry, suggesting that the nature of the chromosomal *rcsD* mutation is contributing to the effect caused by overproduction of the fragments (S3B Fig). An intact Hpt domain in the chromosomal copy of *rcsD* appears to be necessary to allow this modest activation, possibly suggesting that the *rcsD*541 mutant may express a low level of the Hpt domain. This signaling is fully dependent upon RcsB (S3C Fig). Why some of our truncated constructs but not others act in this way is not further investigated here.

### Signaling by RcsD alleles with reduced capacity to interact with IgaA

Plasmids carrying C-terminal portions of RcsD (and thus the Hpt domain) were constructed and tested in both the WT and *rcsD*541 strain. Expression of the ABL-Hpt domains (RcsD_686-C_), the Hpt domain alone (RcsD_792-C_), the cytoplasmic portion of RcsD (RcsD_326-C_), or an *rcsD* allele deleted for the periplasmic region (*rcsD* _Δ45-304_) did not induce signaling in wild type cells (Fig 3B, left panel). However, each of these plasmids led to significantly increased signaling in the *rcsD*541 strain, even in the absence of arabinose (Fig 3B). The cytoplasmic portion of RcsD (RcsD_326-C_), which had no effect on signal in *rcsD*^+^ cells, led to a 12-fold increase in signal in *rcsD541* over the vector control, approximately a 24-fold increase over the normal wild type level of signal (Fig. 3B), grew poorly (6 hour OD_600_ of 0.12, used in Fig 3B), and colonies containing this plasmid became mucoid in the absence of arabinose. Less pronounced induction of signaling was seen in cells expressing the ABL-Hpt domain or the Hpt domain alone in the absence of arabinose. These results are fully consistent with the idea that IgaA represses signaling via interactions with regions in the N-terminus of RcsD; expression of derivatives of RcsD that retain the Hpt domain (and thus can transfer signal from RcsC to RcsB) but that have lost or compromised the IgaA interaction region are now able to activate in the absence of an inducing signal such as PMBN.

We confirmed that this high level of signal is dependent upon RcsC; in an *rcsCD* double mutant, expression of RcsD_326-C_ or RcsD Hpt (*rcsD*_792-C_) does not give rise to signal above the value obtained by adding full length RcsD back to an *rcsCD* double mutant (S3D Fig). The properties of these fragments demonstrate clearly that phosphotransfer from RcsC to RcsB requires, minimally, the Hpt domain of RcsD, and that, in the absence of the IgaA/RcsD inhibitory interaction, RcsC promotes high, constitutive signaling. Finally, because this signaling is not seen in an *rcsD*^+^ strain, signaling by these fragments of RcsD is recessive to the full-length protein.

As noted above, it has been reported that overproduction of the RcsD ABL domain can bind and inhibit RcsB [23]. We further investigated this with overproduction of RcsD regions in our plasmids, testing their potential for induction by PMBN in cells carrying a wild-type chromosomal *rcsD* copy (S3E Fig). If RcsB (or any other step in the pathway) is inhibited, we would expect to block PMBN induction.

Strikingly, while the RcsD plasmid does not affect PMBN-dependent induction in the absence of arabinose (low levels of RcsD), it fully inhibits it when arabinose is present (high levels of RcsD). The plasmid containing the ABL-Hpt domain (Rcs_686-C_) has a very similar profile, as predicted if the ABL domain is necessary for inhibition. RcsD _Δ45-304_, deleted for the periplasmic region, also acts similarly. The Hpt domain itself (RcsD_792-C_) was somewhat less effective in blocking PMBN signaling, consistent with a role for the ABL domain in inhibition. When these plasmids were expressed in a *rcsD*541 host (S3F Fig), RcsD was still able to respond to PMBN without arabinose induction, but not with arabinose. High levels of the ABL-Hpt domain also inhibited the constitutive signaling otherwise seen with low levels of this RcsD fragment, confirming that the inhibition is independent of and downstream of the IgaA/RcsD interaction, not present for this piece of RcsD.

Two of the plasmids gave results that were more difficult to interpret. Expression of a full-length RcsD mutant for the active site histidine (RcsD H842A) had higher activity than expected in an *rcsD*^+^ host but not in the *rcsD*541 host (S3E and S3F Fig). Because this higher activity was seen with or without arabinose, the results suggest that the wild-type RcsD (from the chromosome) and RcsDH842A (from the plasmid) interact, increasing signaling through RcsD. It is also possible that even this low level of RcsDH842A allows some titration of IgaA. The other plasmid that gave unexpected results was RcsD_326-C_. In a wild-type host, this construct acts much like the vector control (low activity without PMBN, increase with PMBN), as if it lacks the inhibitory activity of some of the other constructs. While this might suggest that it is not made in significant amounts, it is clearly able to stimulate activity, independent of PMBN, in an *rcsD*541 host (S3F Fig). It seems likely therefore that for this construct, interaction with RcsD^+^ in the WT host interferes with the ability of the RcsD_326-C_ both to signal (low activity – or + arabinose in *rcsD*^+^ host) and to inhibit (vector-like activity with PMBN, rather than the inhibitory activity seen with ABL-Hpt, containing RcsD_686-C_). These results would suggest that the interaction with the chromosomal RcsD requires sequences upstream of the ABL domain, quite likely the defective HisKA and/or HATPase domains. HisKA domains have been shown to contain dimer interfaces in histidine kinases, and it seems quite possible that the mutant HisKA domain here participates in RcsD/RcsD interactions [25].

These results all suggest that in the absence of arabinose, the pBAD-RcsD constructs express levels of RcsD comparable to the chromosomal level, able to complement and signal but lacking the inhibitory activities seen only with high levels of RcsD achieved after induction with arabinose. To further confirm the behavior of the truncated RcsD proteins, selected alleles were introduced into the bacterial chromosome in place of the native *rcsD* gene. In these strains, RcsD should be expressed from the native promoter, at the native level. These alleles could generally be introduced into an *rcsB* deletion, where signaling is off, but some alleles were difficult to isolate or were clearly unstable in *rcsB*^+^ cells. *rcsD*_326-C_ and *rcsD* _Δ48-304_ could be introduced into the *rcsB*^+^ strain, but cells became quite mucoid and had significant PMBN-independent signaling; RcsD ABL-Hpt was better tolerated (S3G Fig). These results parallel the observations with the RcsD plasmids in an *rcsD*541 strain (Fig 3B; S3F Fig).

Mutant strains were further tested for the ability to support deletion of *igaA*, by P1 transduction from a *bioH*::kan^R^ *igaA*::chl^R^ donor (EAW66, containing a *bioH::kan* mutant closely linked to *igaA*::chl^R^, in a strain containing *rcsD*541), selecting for kanamycin resistance and then testing kanamycin resistant colonies for linkage of the *igaA*::chl^R^ (S3H Fig). In a recipient defective for *rcsB*, rcsC, or *rcsD*, the linkage of the *bioH*::kan and *igaA*::chl^R^ markers is >70%; in a strain WT for the Rcs phosphorelay, linkage was <1 chl^R^/100 kan^R^. Mutations in *rcsD* were introduced into the chromosome, and then used as recipients for P1 transduction (S3H Fig). Strains carrying the *rcsD*541 and *rcsD*841* mutations tolerated loss of *igaA* well, as expected for strains null for *rcsD* (S3H Fig).

*rcsD*_326-C_, carrying all of the cytoplasmic regions of RcsD, did not tolerate loss of IgaA (S3H Fig). Although *chl*^R^ colonies were isolated, those colonies had unstable phenotypes; restreaking yielded colonies that were not as mucoid or fluorescent as the parent strain, strongly suggesting that the *igaA* deletions were only surviving when mutations or deletions of components of the Rcs phosphorelay genes were also present. These results are fully consistent with the behavior of *rcsD*_326-C_ on plasmids (Fig 3B), and consistent with the model that there are critical regulatory contacts between IgaA and RcsD not only in the periplasm but in the cytoplasmic domains as well. Thus, while *rcsD*_326-C_ was negative in the bacterial two-hybrid interaction with IgaA (Fig 2B), the continued dependence on IgaA for viability is consistent with it retaining a critical contact with IgaA. As expected, deleting *rcsB* as well (EAW54, S3H Fig) allowed introduction of the *igaA*::chl^R^ mutation.

Not all *rcsD* alleles could be introduced into the chromosome. *rcsD* _Δ45-304_, which retains a strong interaction with IgaA (Fig 2B) but interferes with cell growth when expressed from a plasmid in *rcsD*541, even without induction (Fig 3B), was lethal, and the chromosomal version of this mutant could not be constructed without accumulating secondary loss-of-function mutations in *rcsD* or *rcsB*. A chromosomal mutant derivative with a slightly longer periplasmic domain deletion, *rcsD* _Δ48-304_, could be constructed, but was mucoid, constitutively active (S3G Fig) and did not tolerate introduction of the *igaA* deletion (EAW106, S3H), consistent with a critical RcsD-IgaA contact that participates in repression beyond the periplasmic region. This allele may be modestly defective for phosphorelay function, and therefore is better tolerated than the *rcsD* _Δ45-304_ allele.

RcsD_686-C_ (ABL-Hpt), which we expect to lack all regions involved in IgaA interaction, led to a lower level of signaling (S3G Fig), is non-mucoid, and, as expected, tolerates loss of *igaA* (EAW108, S3H Fig). We would suggest, based on its phenotypes, that this construct is not fully active for passing signal from RcsC to RcsB.

These results lead to the following conclusions: 1) the RcsD periplasmic region is essential for full interaction with IgaA and, most strikingly, for full inhibition by IgaA, but is not sufficient for binding or titration of IgaA. This is most consistent with a direct interaction of the RcsD periplasmic loop and IgaA. The precise role of the trans-membrane (TM) regions flanking the periplasmic loop have not yet been explored. 2) An additional region or regions of interaction exist, in the cytoplasmic PAS-like domain of RcsD; this interaction is not sufficient to allow IgaA-dependent repression, but presumably improves binding in the presence of the TM/periplasmic region (thus allowing binding to and titration of IgaA) and contributes significantly to repression by IgaA. 3) Constructs with the Hpt domain but lacking the periplasmic loop (or lacking both the periplasmic region and TM helices) of RcsD are capable of Rcs induction-independent signaling, presumably because they are blind to IgaA repression. 4) The Hpt domain on its own, or the full cytoplasmic domain, is recessive to RcsD^+^, and is thus not able to constitutively signal in the presence of functional RcsD.

### Critical residues in the cytoplasmic PAS-like domain of RcsD

Alanine scanning mutagenesis of individual conserved residues in the PAS domain region of RcsD was carried out in the pBAD-RcsD plasmid. Plasmids were initially screened for level of fluorescence in an *rcsD*541 mutant strain, in the absence of arabinose. In this assay, functional RcsD expressed from the plasmid reduces fluorescence by complementing the *rcsD*541 allele, nulls would be expected to not affect fluorescence, and mutations that were capable of passing phosphate from RcsC to RcsB but were less sensitive to IgaA repression might be expected to have higher fluorescence (see right-hand panel in Fig. 3B, for example). Unexpectedly, many of the plasmids that appeared to give strong signals and were thus thought to possibly have become blind to IgaA instead had acquired stop codons. These mutants were not further investigated here. Thus, we instead focused on alanine mutants that retained function, measured by the ability to complement the *rcsD*541 mutant, reducing the elevated signal found in this mutant to the lower level found in *rcsD*^+^ strains (compare lane 1 and lane 3, S4A Fig). Among the six mutants screened, one was striking in that it was unresponsive to PMBN induction, suggesting that it somehow locked RcsD into an “off” configuration. This mutant allele, RcsD T411A, was further analyzed.

The *rcsD* T411A mutation was introduced into the chromosome and tested for its response to PMBN (Fig 4A). The mutant had a lower basal level of Rcs signaling and, as was seen with the plasmid-borne copy, this mutant had a very muted response to PMBN. A22, an inhibitor of MreB, and mecillinam have also been reported to induce the Rcs System [3, 26], and we confirmed that induction with our reporter (Fig 4A, WT). The T411A mutant also failed to respond to A22 or mecillinam (Fig 4A, red bars).

**Figure 4:**
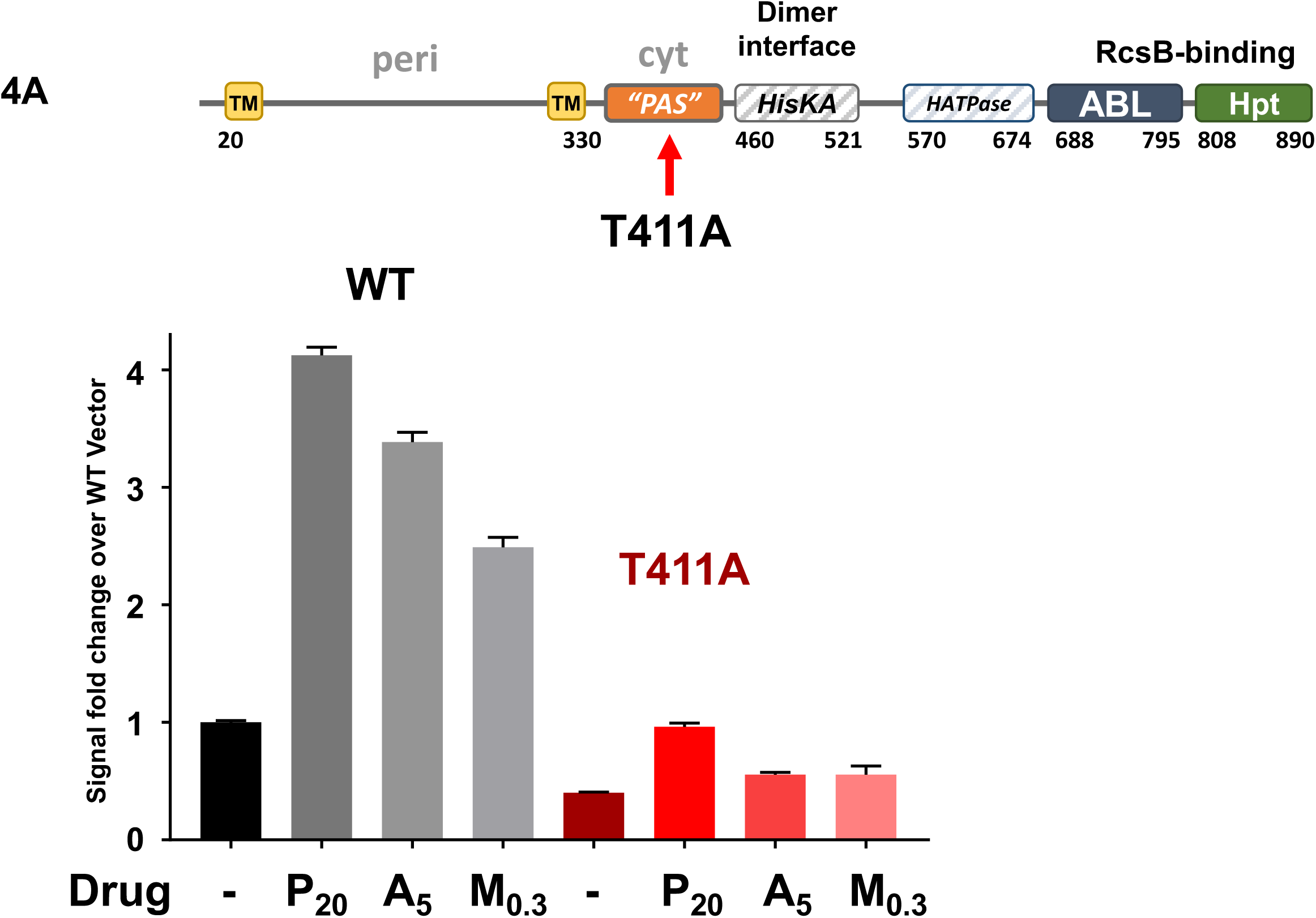
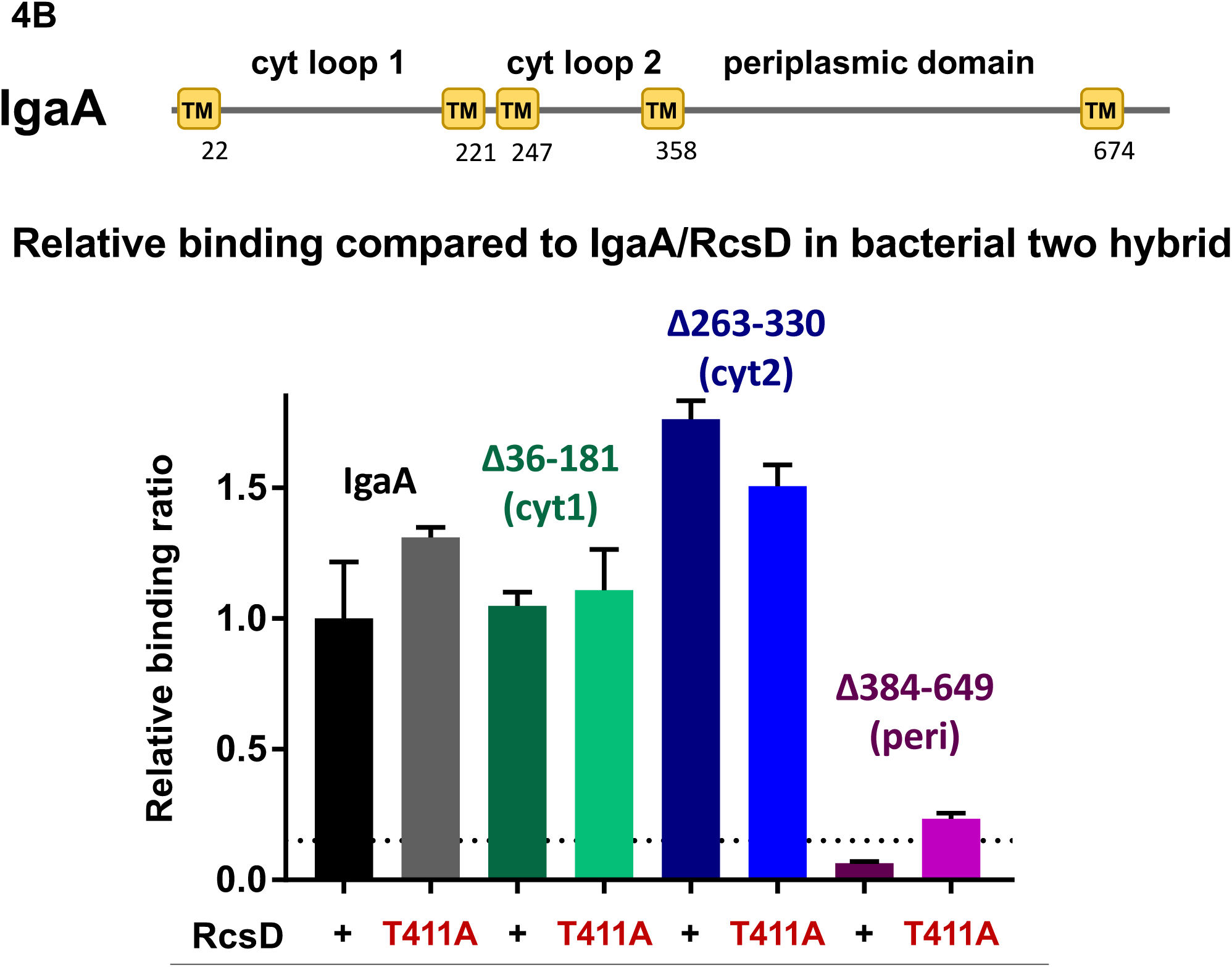
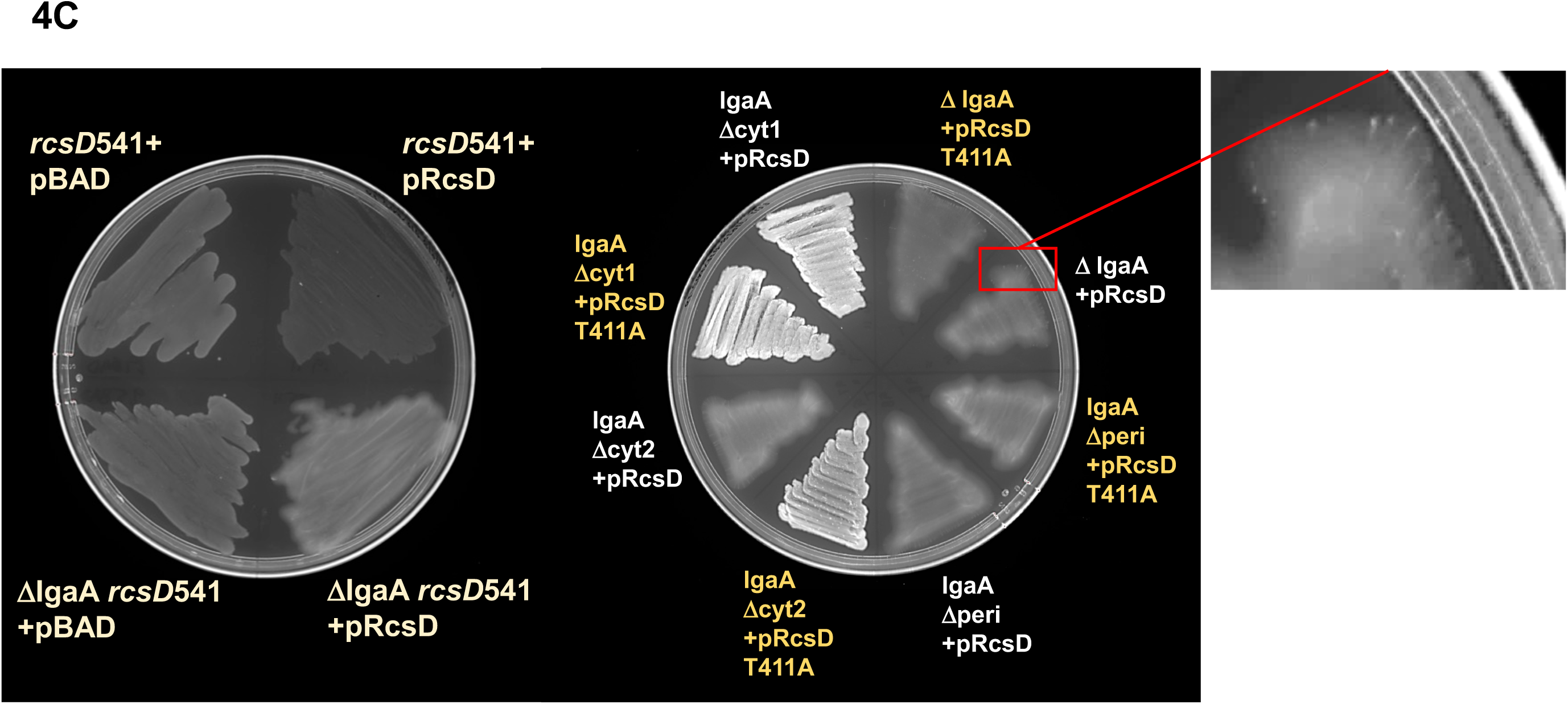
An RcsD mutation that blocks Rcs induction by increasing IgaA interaction. A. RcsDT411A does not respond to Rcs stimuli. Both wild type and *rcsD*T411A strains (EAW8 and EAW121) were treated with nothing (-), 20µg/mL polymyxin B nonapeptide (P_20_), 5µg/mL MreB inhibitor A22 (A_5_) or 0.3µg/mL Mecillinam (M_0.3_). Both A22 and Mecillinam give a lower dynamic range of wild type signaling than PMBN. B. RcsD “PAS” domain mutation T411A seems to confer tighter binding to an IgaA construct that is missing the periplasmic domain. IgaA schematic includes yellow transmembrane domains (TM), amino acid numbering, and loops labeled with their localization. BACTH results are shown as ratios relative to the wild type IgaA/RcsD interaction. Wild-type RcsD and RcsD T411A are comparable in binding IgaA constructs containing the main periplasmic loop; RcsD T411A interacts with IgaA deleted of its periplasmic loop at a detectable level, not seen with WT RcsD. Plasmids, background controls and fold above background values are shown in S4B Fig. C: RcsD and RcsD T411A show different levels of Rcs dysregulation when introduced on plasmids into cells carrying the *rcsD*541 mutation and chromosomal *igaA* deletions. The left plate contains (clockwise from top left quadrant) *rcsD*541 with vector (EAW19 with pBAD24), EAW19 containing RcsD on a plasmid (pEAW11), *rcsD*541 in a complete *igaA* deletion (EAW95) with vector, and EAW95 containing pEAW11. Evident in this panel is 1) a decrease in fluorescence when RcsD is complemented from EAW19 (evident on graphs in Fig 3A, 3B), 2) mucoidy in EAW95 when RcsD is complemented, with only slightly raised apparent fluorescence. Mucoidy scatters the mCherry fluorescence, making it appear lower than the actual output per cell. Right panel and inset: *igaA* deletions in the chromosome in an *rcsD*541 background show mucoidy, signaling, and instability upon introduction of plasmids containing RcsD or RcsD T411A. The inset shows bright streaks within EAW95+pEAW11; this mucoid primary transformant spontaneously generates non-mucoid *rcs* mutants. Because many mutants are not nulls, and the loss of mucoidy increases the apparent fluorescence, these show up as more brightly fluorescent, even though their level of P_rprA_-mCherry signal is lower. On the right plate RcsD and RcsD T411A plasmids lead to high levels of mucoidy in either a strain carrying a full *igaA* deletion or a strain carrying a deletion of IgaA periplasmic loop (EAW95, EAW96). Cells carrying a deletion of cytoplasmic loop one (EAW98) and either the RcsD or RcsD T411A plasmids are highly fluorescent but not mucoid, suggesting a somewhat less critical role for this loop in mediating IgaA repression. Only deletion of IgaA cytoplasmic loop two deletion shows a significant difference dependent on the RcsD allele; lower levels of Rcs activation were seen (loss of mucoidy) in the presence of RcsDT411A (EAW97 with pEAW11T).

We can imagine two general ways in which T411A might block induction. It might affect the ability of signal to move through the phosphorelay, possibly locking RcsD in the “phosphatase” confirmation. In this case, we might expect it to be indifferent to the presence of IgaA. In the alternative model, *rcsD*T411A is locked off because it no longer releases the interaction with IgaA when signal is received; if so, it will still be sensitive to loss of IgaA. To test whether the RcsD T411A was causing a “locked” state in which the protein could no longer pass signal to RcsB, we tried to delete *igaA* in the chromosome. The strain did not tolerate IgaA deletion (EAW121, S3H Fig), which suggested that RcsD T411A abrogates activation by increased or changed interaction with IgaA.

We next turned to IgaA to begin to identify the regions likely to interact with RcsD. Based on our observations with the RcsD truncations, we would predict at least two regions of interaction betweenRcsD and IgaA: between the RcsD periplasmic region (critical for IgaA regulation) and the periplasmic region of IgaA, as well as additional important interactions between the cytoplasmic PAS-like region of RcsD (and possibly other regions) and cytoplasmic domains of IgaA.

The periplasmic domain of IgaA (see Fig 4B) has previously been found to interact with RcsF [8]. Here, we find that deletion of the periplasmic domain (IgaA _Δ384-649_) fully abrogates the interaction of RcsD and IgaA; T411A modestly restores this interaction, consistent with the mutant leading to increased interaction within the cytoplasmic regions (Fig 4B and S4B Fig). Deletion of either cytoplasmic loop 1 or cytoplasmic loop 2 of IgaA had essentially no effect on the interaction with wild-type RcsD, suggesting that the primary interactions that drive the bacterial two hybrid signal are between the periplasmic regions. Periplasmic point mutation L643P is a stable protein (S4C Fig) that caused a partial loss of function mutant in *igaA* [27]; this mutation led to loss of interaction of IgaA and RcsD (S4D Fig). However, other alleles at this position (L643A) or nearby did not disrupt interaction or activity (S4D and S4E Fig), suggesting that L643 is not itself a critical residue but that L643P may disrupt IgaA folding or localization. The specific regions within the IgaA periplasmic domain that contact RcsD remains for future analysis.

Chromosomal deletions of either one of the *igaA* cytoplasmic loops (Δ36-181, cyt1; Δ263-330, cyt2) or of the periplasmic loop (Δ384-649, peri) (see Fig 4B) were constructed in cells mutant for *rcsD* and carrying the P_rprA_-mCherry reporter. A complete *igaA* deletion was used for comparison. Introduction of the RcsD plasmid, even in the presence of glucose (low levels of RcsD expressed) was poorly tolerated in all the *igaA* deletions, with secondary mutations arising at a rapid rate (see inset, Fig 4C). Therefore, assays in liquid were considered untrustworthy, and the phenotypes of the primary transformants were evaluated on agar plates (Fig 4C).

Transformation of the RcsD plasmid into cells carrying a deletion of *igaA*, the *igaA* periplasmic domain (*igaA* _Δ384-649_) or the second cytoplasmic domain (*igaA* _Δ263-330_) gave rise to highly mucoid growth, consistent with lack of IgaA function. Introduction of the RcsD^+^ plasmid into cells deleted for the first cytoplasmic domain (*igaA* _Δ36-181_) gave less mucoid growth, although the P_rprA_-mCherry reporter was well expressed compared to WT and *rcsD*541 (Fig 4C left panel), consistent with an important role for this domain of IgaA as well. The plasmid expressing RcsDT411A rather than RcsD^+^ was introduced into these strains. This mutation was unable to decrease the signal in the full deletion of *igaA* or the deletion of the periplasmic domain. However, it reduced mucoidy and allowed more robust colony growth in cells carrying the second cytoplasmic domain deletion (Fig 4C). This result is most consistent with RcsDT411A improving the interaction with IgaA cytoplasmic loop 1 and therefore abrogating induction (Fig 4A).

### Analysis of RcsC domains and involvement in signaling

From previous work, it is clear that RcsC plays an essential role in signaling in the Rcs phosphorelay, as the source of phosphorylation [16]. However, as shown above, full length RcsC did not interact with IgaA in the bacterial two hybrid assay. The plasmids expressing T18 and T25 fusions to full-length RcsC interfered with cell growth, and also did not interact with RcsD, and thus while our data strongly supports the interaction of IgaA with RcsD, we are cautious in interpreting this negative result with RcsC and IgaA. A construct expressing only the cytoplasmic domains of RcsC (rcsC_326-C_) interacts well with RcsD as well as with the cytoplasmic regions of RcsD, although a bit less strongly (S5A Fig). It interacts as well with a version of RcsD missing residues beyond aa 683 (RcsD_N-683_), but not at all with RcsD_N-525_, suggesting that the region between aa 525 and 683 of RcsD, including its inactive HATPase domain is essential for this interaction (S5A Fig). Given that phosphate flows from the C-terminal REC domain of RcsC to the RcsD Hpt domain, we would predict a further, possibly transient, interaction between the RcsC REC domain and the RcsD Hpt domain.

Unlike RcsD, RcsC constructs in pBAD24 were often cytotoxic, causing massive cell lysis without any detectable increase in P_rprA_-mCherry signal above background, and slow growth even in rich defined glucose media, where the pBAD promoter should be only modestly active. To avoid this overproduction toxicity, deletions and substitutions of interest were introduced into the chromosomal copy of *rcsC* and tested for response to induction by PMBN. RcsC carrying a mutation in the kinase active site (H479A) had low activity and was not responsive to PMBN, as expected (Fig 5A). Note that the activity in this mutant is more like the wild-type strain without PMBN than like the deletion of *rcsC* (Fig 5A), in support of experiments reported by Clarke et al that the active site His is not necessary for phosphatase activity [16].

**Figure 5:**
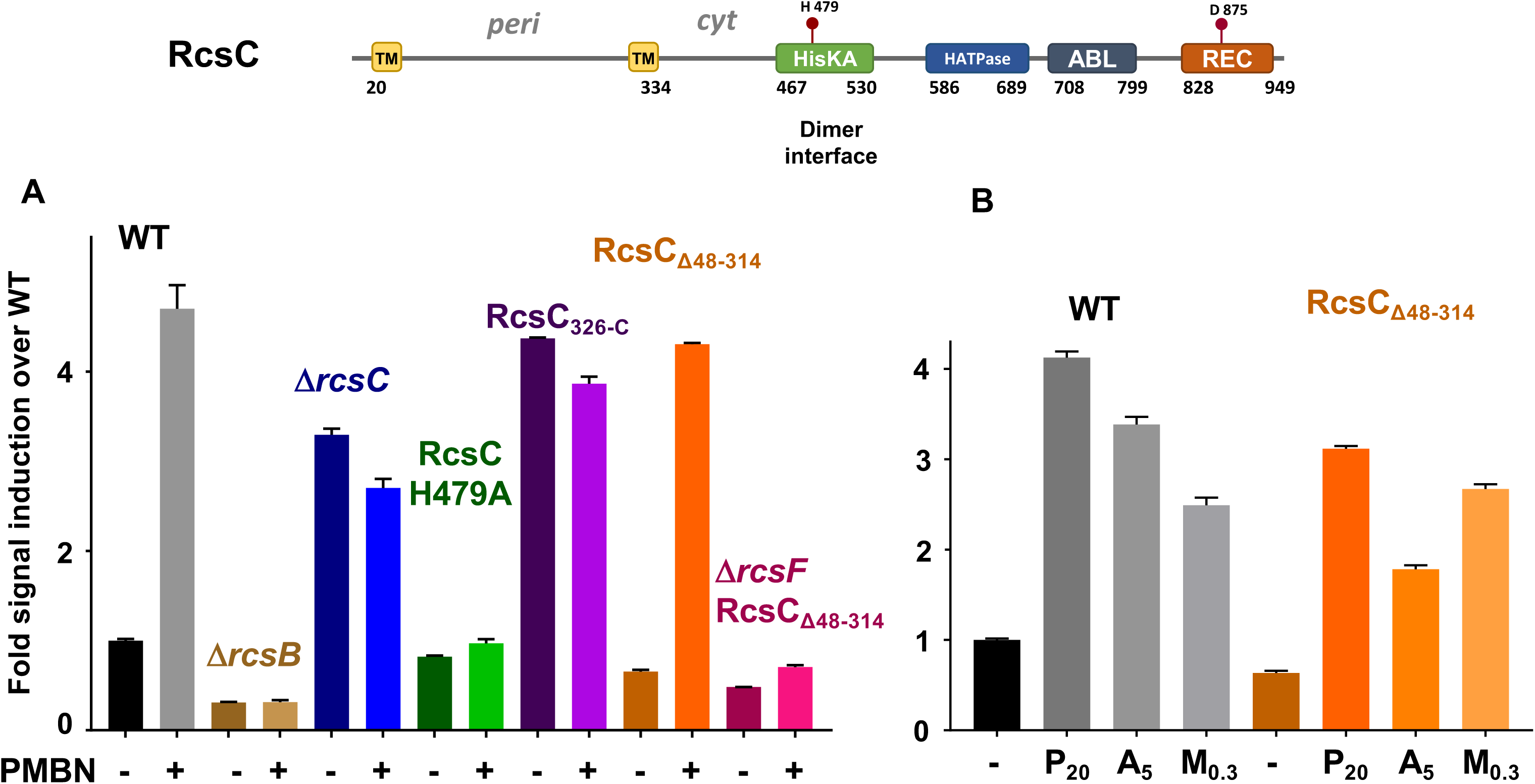
The RcsC periplasmic region is unnecessary for polymyxin B nonapeptide (PMBN), A22, and mecillinam-induced signaling. The top panel shows a schematic of RcsC with domains, topology and active site residues noted. A. PMBN induction in various *rcsC* mutations. Strains included are (L to R) EAW8, EAW31, EAW91, EAW92, EAW56, EAW70 and EAW85. B. The effect of three Rcs stimulating drugs, PMBN, A22 and mecillinam (P_20_, A_5_, M_0.3_) on WT and RcsC_Δ48-314_. The RcsC periplasmic deletion strain has a lower basal level of signal than wild type but shows large increases in signal when exposed to Rcs-inducing drugs.

Unexpectedly, a deletion of the periplasmic portion of RcsC, leaving the TM helices (RcsC_Δ48-314_), had a lower basal level of signal than WT, yet responded strongly to PMBN and A22 (Fig 5 A, B). This allele still requires RcsF for PMBN signal detection, which suggests that this signal comes from RcsF through IgaA to RcsD to RcsC (Fig 5A). Cells carrying the *rcsC*_Δ48-314_ mutation tolerate IgaA deletion, although they become mucoid and unstable (EAW70, S3H Fig); deletion of *rcsC* or *rcsC*H479A was unaffected by loss of *igaA* (S3H Fig). We suggest that this allele has a modestly decreased ability to signal, which in other experiments allows cells to support deletion of *igaA*. Overall, this result strongly suggests that IgaA regulation of the phosphorelay and signal transduction via RcsF are not acting through the periplasmic loop of RcsC.

Although the periplasmic region is not necessary for RcsC function, it would appear that membrane association is. Cells carrying a deletion of the membrane spanning portion (RcsC_326-C_) acted in a similar manner to an *rcsC* deletion, with a constitutive level of reporter expression and no response to PMBN (Fig 5A). Consistent with a loss of function for the rcsC_326-C_ allele, the deletion of *igaA* could be introduced into this strain, and cells remained non-mucoid (EAW56, S3H Fig). A chimeric construct in which the MalF TM and periplasmic region replaced the *rcsC* periplasmic region restored the ability of the cell to respond to PMBN (S5B Fig), albeit with a higher basal level of signaling in the absence of PMBN. Finally, a series of periplasmic deletions with different linker lengths all responded to PMBN to some extent, although constructs with shorter turns had somewhat lower basal levels (S5B Fig).

Although these results demonstrate that the RcsC periplasmic region is not necessary for sensing the OM stress signal elicited by PMBN, it seemed possible that other inducing stresses, such as mecillinam or A22, might act in a way that was dependent upon the periplasmic region of RcsC. This was tested in our system and demonstrated that the RcsC_Δ48-314_ mutation still showed induction in response to all three stimuli (Fig 5B). These additional Rcs stimuli elicited an increase in P_rprA_-mCherry that had less dynamic range and more cell death than PMBN. At the published concentrations, A22 (5 µg/mL; Sigma) and Mecillinam (0.3 µg/mL; Sigma) could induce the WT strain and the RcsC _Δ48-314_ strain, while RcsDT411A failed to respond (Fig 4A). Therefore, for A22 and Mecillinam, as for PMBN, the *rcsC* periplasmic region is not required for sensing and responding to signal.

## DISCUSSION

The results reported here provide a new view of how IgaA transduces inducing signals within the complex Rcs phosphorelay (Fig 1A). IgaA, a multipass membrane protein, is a strong negative regulator of Rcs. As previously described, signals, such as PMBN, believed to disrupt LPS interactions, or A22 and Mecillinam, peptidoglycan disruptors, change the nature of the RcsF/IgaA interaction. This leads to a change, presumably a decrease, in IgaA’s interaction with the downstream phosphorelay. We find that the point of interaction of IgaA is with the phosphotransfer protein RcsD, rather than with the RcsC histidine kinase. In fact, while RcsC function requires membrane association, the TM and periplasmic sequences of RcsC are not required (Fig. 5, S5B Fig). The change in the IgaA-RcsD interaction frees RcsC-generated phosphate to flow from RcsC to RcsD, and from there to RcsB, activating signaling downstream of RcsB. Deletion and mutation analysis of RcsD identified multiple regions important for IgaA-dependent regulation, separate from the regions critical for phosphorelay signal flow from RcsC to RcsD. These observations help to explain why RcsD includes not only an Hpt domain but also trans-membrane and signaling domains. We suggest that the use of RcsD as the direct target of IgaA has allowed the development of a poised signaling complex, without impinging on structures necessary for histidine kinase activity. In addition, this branched signaling pathway allows the possibility of other signals directly regulating RcsC activity.

### Multipartite interactions of RcsD and IgaA regulate signaling: anchors and switches

Our analysis of the regions of RcsD and IgaA necessary for interaction and regulation suggest multiple points of contact between these proteins, each with different roles in regulation. The first contact is in the periplasmic loops of these two proteins. The periplasmic domain of RcsD (amino acids 45-304) is necessary but not sufficient for repression by IgaA; the periplasmic region of IgaA (aa 384-649) is similarly essential for IgaA function and drives the interaction of IgaA and RcsD in a bacterial two-hybrid assay (Fig 2B, Fig 4B). We assume that these periplasmic domains directly contact each other (Fig 6). Because others have demonstrated that overproduction of IgaA missing the periplasmic domain can support repression and allow depletion of wild-type IgaA [8], we suggest that this strong periplasmic interaction provides an anchor for interaction with RcsD, but likely not the region in which signal is sensed. Thus, the periplasmic contact can be bypassed by overproduction, but is critical for repression at normal levels of the interacting proteins.

**Figure 6:**
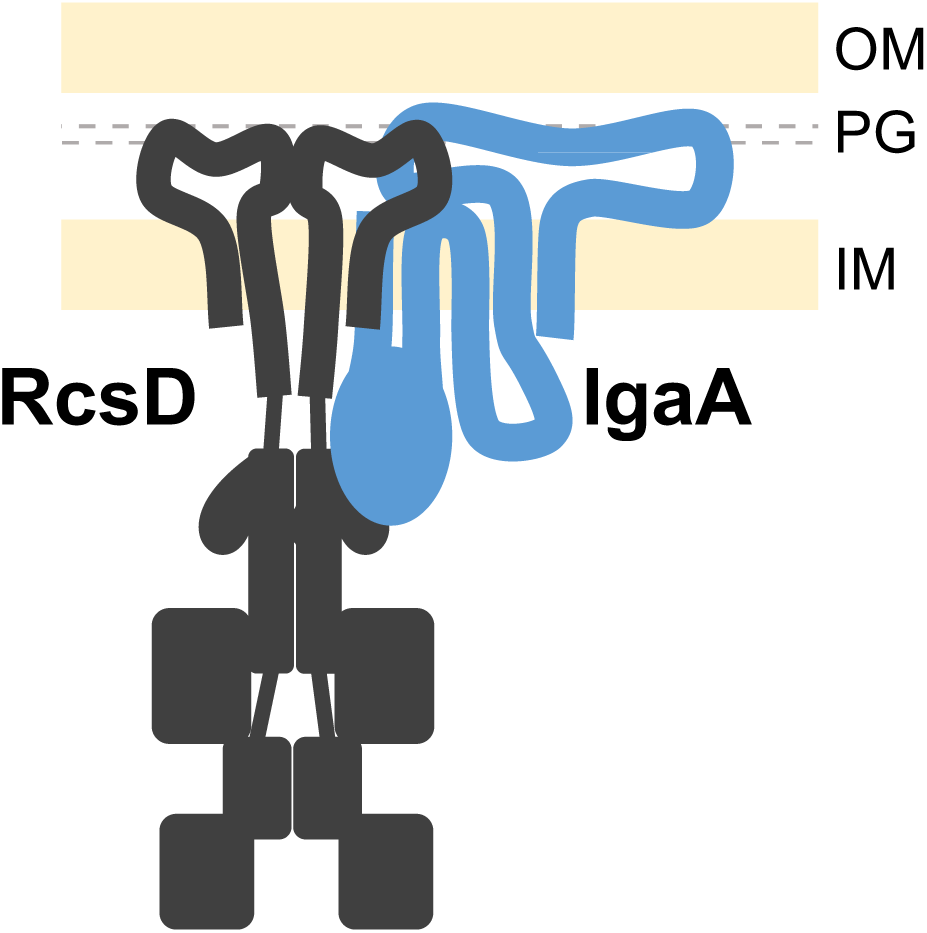
Proposed interactions of IgaA and RcsD. Extensive interactions in the periplasm and in the cytoplasm are shown, consistent with genetic data indicating that signals pass from one compartment to the other via IgaA. Anchoring interactions in the periplasm drive the BACTH interaction signal, and are required for IgaA repression of signaling. Interaction of the IgaA cytoplasmic loop 1 (blue oval) and the “PAS” domain of RcsD are suggested to comprise the signaling interaction, tightened in *rcsD*T411A, an allele that blocks induction.

The cytoplasmic PAS-like domain of RcsD is also necessary for regulation, interaction in the bacterial two-hybrid assay and interaction as judged by titration of IgaA (Fig 2B, Fig 3A). It seems likely that this cytoplasmic domain contacts one or both cytoplasmic loops of IgaA. Both cytoplasmic loop 1 and cytoplasmic loop 2 of IgaA are necessary for RcsD to function properly (Fig 4C), in agreement with previous work [8]. Because the bacterial two-hybrid interactions are primarily driven by the periplasmic domains (Fig 4B), the contacts of RcsD with the IgaA cytoplasmic loops are likely to be weaker. We do not currently have any direct evidence that cytoplasmic loop 2 is contacting RcsD, but certainly deletion of this loop, like deletion of the periplasmic region, abrogates repression (Fig 4C). We suggest that the interaction of cytoplasmic loop 1 and RcsD, in the region around T411, constitutes the regulatory switch for this system. Deletion of loop 1 is the least detrimental in terms of bacterial growth and signaling (Fig 4C), suggesting that the contacts outside cytoplasmic loop 1 are sufficient for enough IgaA repression of RcsD to support viability. Our model suggests that the additional repressive interaction in loop 1 is normally lost upon Rcs stimulus (in the presence of PMBN, for instance), and that the anchor contacts in the periplasm and with IgaA loop 2 ensure that signaling is never so high that the cell dies. In the T411A mutant, this stimulus-sensitive contact becomes stronger, so that the system becomes uninducible (Fig 4A). This can be seen in the bacterial two-hybrid assay as some restored interaction in the absence of the IgaA periplasmic region (Fig 4B).

In work by Collet and coworkers, overproduction of cytoplasmic loop 1 [8] was, by itself, capable of repressing an Rcs reporter to a similar extent to that seen with both cytoplasmic regions, further supporting a critical role for this region of Rcs.

### RcsD is an unusual phosphorelay protein

Phosphate flow in complex phosphorelays such as Rcs is from His (kinase domain) to Asp (response regulator domain of RcsC) to His (RcsD phosphotransfer protein) to Asp (RcsB response regulator domain). RcsD, is a large inner membrane protein with many additional domains; its domain organization suggests that duplication of an ancestral protein may have given rise to RcsC and RcsD. Our results suggest critical roles for these additional regions of RcsD.

Consistent with its role as an anchor for IgaA, alignments suggest significant regions of conservation within the periplasmic domain of RcsD, apparently more so than the similarly sized RcsC periplasmic domain, which we show here is not critical for signaling (Fig 5A, S6 Fig). There is significant conservation as well in the truncated PAS domain, but less conservation in the inactive HATPase domain than in the active parallel RcsC domains. Future work will be necessary to identify the periplasmic interaction points of RcsD with IgaA and to understand whether the RcsD ATPase domain plays any critical role in regulation.

### Alternative signaling pathways remain to be understood

The complexity of the Rcs phosphorelay provides opportunities for signals to regulate RcsB activity independently of the RcsF-IgaA-RcsD interaction network. Some transcription factors interact directly with RcsB, independent of its phosphorylation, to make heterodimers that regulate specific sets of genes (reviewed in [1]). In addition, there is evidence for activation of RcsB-dependent genes, dependent upon RcsC and RcsD, but independent of RcsF. The two cases in which this has been reported involve overproduction of the DjlA DnaJ-like chaperone and mutation in the periplasmic disulfide bond formation protein, DsbA, possibly suggesting that alterations in protein folding may be the inducing event [1, 16, 28].

One other unexplored aspect of our work is the possible expression of low levels of the C-terminal domains of RcsD, to produce a short phosphotransfer protein that would not be subject to IgaA regulation. For instance, the modest activity of RcsD_N-522_ and RcsD_N-683_ in the *rcsD*541 host (S3A Fig) was unexpected. Because this same increase was not seen when the host contained other *rcsD* alleles (*rcsD*841*, for instance), we suggest it may be due to low level expression of a C-terminal fragment of the chromosomal RcsD protein able to transfer phosphate from RcsC to RcsB. In other experiments, we found that unplanned stop codons were found in some plasmids expressing *rcsD* alanine scan mutants. These plasmids, rather than acting like nulls, had activity significantly above that of a null strain, again suggesting that they might be expressing a C-terminal fragment of RcsD. Whether this is ever made under wild-type physiologically relevant conditions remains to be determined but would provide the possibility of an IgaA-resistant signaling pathway.

Overall, while a critical step in the best understood signaling pathway is clarified here, there is still much to learn about Rcs, other modes of signaling to the phosphorelay, and exactly how the IgaA/RcsD interactions modulate phosphate movement from RcsC through RcsD to RcsB. Given the range of genes regulated by RcsB, and the importance of these genes for bacterial behavior, the options for multiple ways for the system to be regulated may not be surprising.

## Materials and Methods

### Bacterial growth conditions and strain construction

Cells were grown in LB with appropriate antibiotics (ampicillin 100µg/mL, kanamycin 30-50µg/mL, chloramphenicol 10µg/mL for the *cat* cassette in *cat sacB* strains and 25µg/mL for others (*chl*^R^), tetracycline 25µg/mL, gentamicin 10µg/mL, zeocin 50µg/mL); 1% glucose was added in some cases to reduce basal level expression of P_BAD_ and P_Lac_ promoters. For fluorescence/growth assays, strains were grown in MOPS minimal glucose or minimal glycerol (Teknova). Strains were constructed via recombineering and/or P1 transduction with selectable markers, as outlined in S1 Table. Strains, plasmids,oligonucleotides and gBlocks used in this study are listed in S1-S3 Tables. Oligonucleotides and gBlocks were from IDT DNA, Coralville, IA.

For recombineering, cells carrying the chromosomal mini-*λ* Red system or the plasmid-borne Red system (pSIM27) were grown in LB, without or with Tetracycline respectively, at 32°C to an OD_600_ of ∼0.4-06. At mid-log, cultures were transferred to a water bath at 42°C to induce expression of the *λ*-Red system for 15 minutes and then immediately chilled in an ice-water slurry for 10 minutes prior to washing in sterile ice-cold water to make electrocompetent cells. 100 ng of ss oligo DNA or dsDNA (PCR product or gBlock) were used in the electroporation; 1 ml of LB or SOC was added for recovery before plating on selective plates [29]. Truncations and point mutations were introduced in place of the wild-type chromosomal copies of genes, leaving no marker or scar, unless otherwise indicated, by first inserting the counter-selectable ara-kan-kid cassette from CAI_91 and simultaneously deleting the gene of interest, and then replacing it with the desired allele, provided either as a PCR product or a gBlock. This cassette, a gift of C. Ranquet (BGene Genetics, Grenoble), expresses the Kid toxin under the control of an arabinose-inducible promoter. Cells carrying the *ara*-*kan*-*kid* counter-selectable marker cassette were grown with added 1% glucose in the media to repress. Counter-selection for removal of the *ara*-*kan*-*kid* cassette was done on LB-1% arabinose plates. All plasmid and chromosomal mutations were confirmed by sequencing using flanking primers.

### DNA and strain manipulation and mutagenesis

Polymerase chain reactions were performed using Pfu Ultra II polymerase (Agilent) or Clontech Hifi polymerase (Takara). Primers used in this study are listed in S3 Table. PCR products were purified using column purification (Qiagen) according to the manufacturer’s instructions. Gibson assemblies were performed using the Clontech In-Fusion HD Cloning Kit (Takara) and transformed into either Clontech Stellar Cells or NEB Turbo cells containing LacI^q^.

Alanine-scanning mutagenesis was carried out by SGI-DNA (San Diego, CA), using their BioXP system, on pBAD24-RcsD (pEAW11). We ordered single mutants targeted to conserved residues within the cytoplasmic region of RcsD, from residue 326-683. Plasmids were first transformed into Stellar *E. coli* (Clontech), extracted and retransformed into EAW19, screening for fluorescence on minimal glucose-ampicillin agar plates, in comparison to cells carrying pBAD-RcsD^+^ or the empty pBAD vector. Out of 35 mutants screened, ten had fluorescence levels comparable to the pBAD-RcsD^+^ control; six were further studied (S4A Fig). Another 17 had higher fluorescence than either the pBAD vector or the pBAD-RcsD^+^ control, but sequencing of these isolates showed that they had all contained, in addition to the designed mutation, additional unexpected stop codon mutations and were not further studied here.

### Bacterial Adenylate Cyclase Two-Hybrid Assay

In the bacterial adenylate cyclase two hybrid assay (BACTH), an adenylate cyclase mutant strain is used to assay for beta-galactosidase activity engendered when the T18 and T25 portions of adenylate cyclase are reconstituted, allowing cAMP/CRP to activate the *lac* operon. On their own, T18 and T25 will not form adenylate cyclase efficiently unless they are fused to two interacting proteins [21]. Tags were C-terminal to avoid interference with protein insertion into the membrane.

The RcsD and RcsC fusion proteins were tested for determine if they were functional and thus presumably membrane-localized. Plasmids expressing RcsD-T25 and RcsC-T25 were introduced into strains containing deletions for those two genes; after transformation, the cells were transduced with P1 grown on a strain NM357, containing *igaA*::chl^R^, selecting for chloramphenicol resistance. In a strain deleted for *rcsD* or *rcsC*, the *igaA* deletion can be introduced by P1 transduction. However, the fusion plasmids blocked the ability of cells to be transduced with *igaA*::chl^R^, consistent with them complementing their respective deletions (S2E Fig).

### *igaA* co-transduction frequencies

*bioH*/*igaA* co-transduction frequencies were used to determine which strains could support loss of IgaA. *bioH*, at 3544844 nt, is linked to *igaA* (position 3526469). The *bioH*::kan^R^ mutant from the Keio collection [30] was introduced by P1 transduction into an *rcsD*541 *igaA*::chl^R^ mutant (EAW17), selecting for kanamycin resistance and retention of chloramphenicol resistance (*igaA*::chl^R^), to create EAW66. Because *rcsD* is inactive in this strain, it can tolerate loss of *igaA*. P1 transduction from this donor to recipient strains was carried out, selecting for Kanamycin Resistance and then screening 50-100 colonies for linkage to *igaA*::chl^R^. In *rcsB*, *rcsC* or *rcsD* null recipients, the co-transduction frequency was 78%. In a wild-type strain, the linkage dropped to zero, consistent with the known lethality of an *igaA* deletion [3, 31] (S3H Fig).

### Fluorescence assays

Fluorescence assays for Rcs activation were performed in 96 well plates in a Tecan Spark 10m spectrophotometer. These strains carried a transcriptional fusion of mCherry, at the *ara* locus, to the promoter for sRNA RprA, as a reporter for Rcs pathway activation, referred to here as P_rprA_-mCherry. Fluorescence of cells was measured in MOPS glucose minimal media (Teknova) unless otherwise stated. The pBAD24 plasmid was used for overexpression of RcsD fragments in strains expressing *araE* constitutively to ensure homogenous arabinose uptake [32]. For cells expressing proteins from pBAD, overnight cultures in MOPS minimal glucose were washed with MOPS minimal glycerol to eliminate residual glucose, then diluted into fresh MOPS minimal glycerol media (.05% glucose, .5% glycerol) with 0.02% arabinose or 0.2% glucose as an uninduced control. Polymyxin B nonapeptide (PMBN; Sigma), a non-toxic polymyxin derivative, was used at 20 ug/mL to produce Rcs induction. To check for Rcs induction by other known compounds, A22, an MreB inhibitor was used at 5ug/mL, and mecillinam was used at 0.3ug/mL.

Each strain/condition combination was performed in technical triplicate on the plate, with biological replicates performed on different days. Optical density and mCherry fluorescence were monitored every fifteen minutes for seven hours. At the end of six hours, measurements of fluorescence at equivalent OD_600_ values (0.4 +/- 0.03 after starting at OD_600_ .03-.05) were converted to bar graphs of fold change of fluorescence with respect to the wild type strain. Some strains arrested growth early and never achieved 0.4 OD_600_, and the OD_600_ at 6 hours for those are noted on the graph. Six hours marks the time when the wild type strain begins to transition to stationary phase, and ODs become less interpretable due to cell aggregation in the well bottom.

## Acknowledgments

We thank S. Buchanan and members of her laboratory for discussion throughout this work, as well as members of the LMB. Erin Wall was supported during much of this work by a PRAT Fi2 fellowship GM123943 from NIGMS. This research was supported in part by the Intramural Research Program of the NIH, National Cancer Institute, Center for Cancer Research.

## Supporting Information

**S1 Fig.**
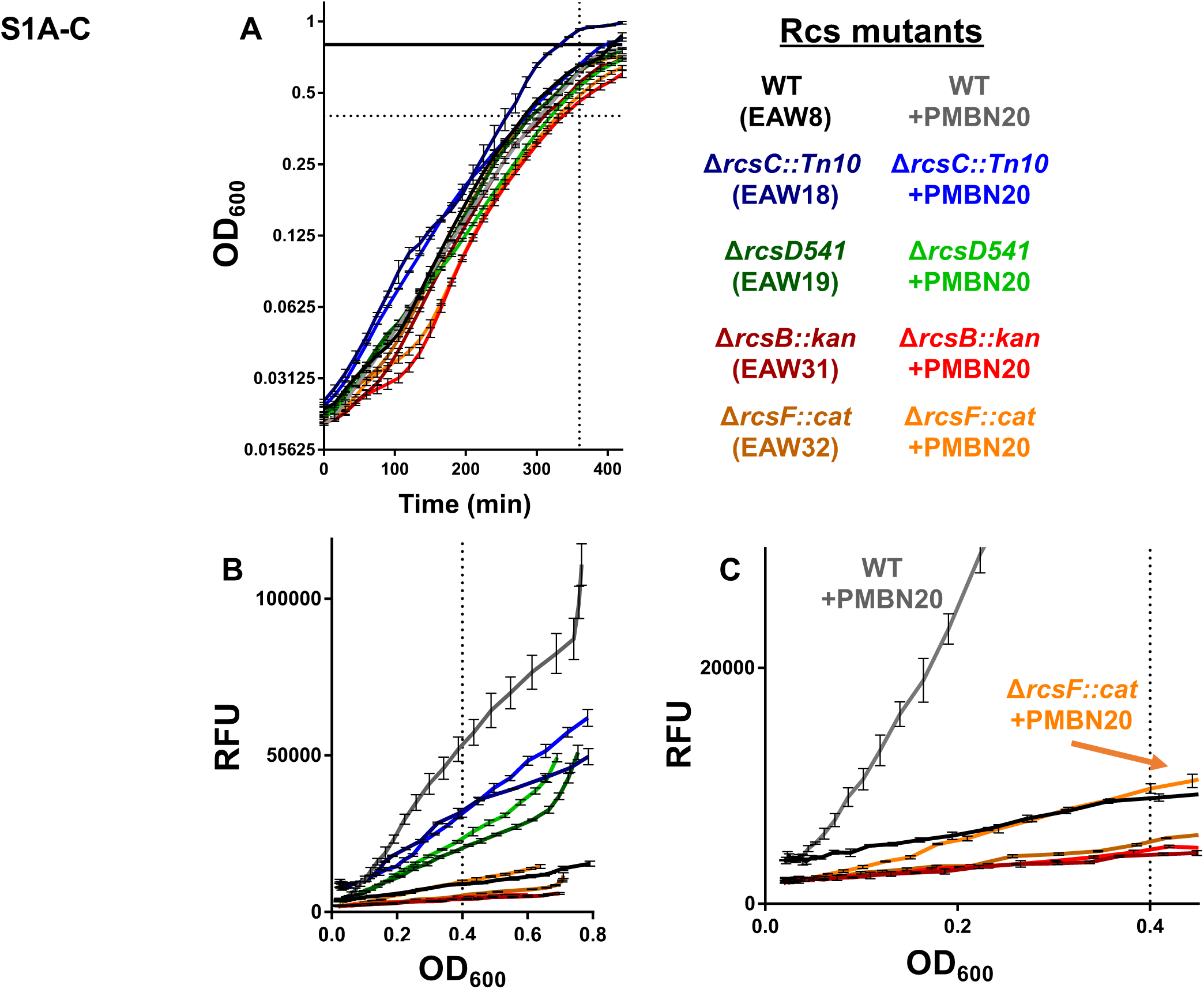
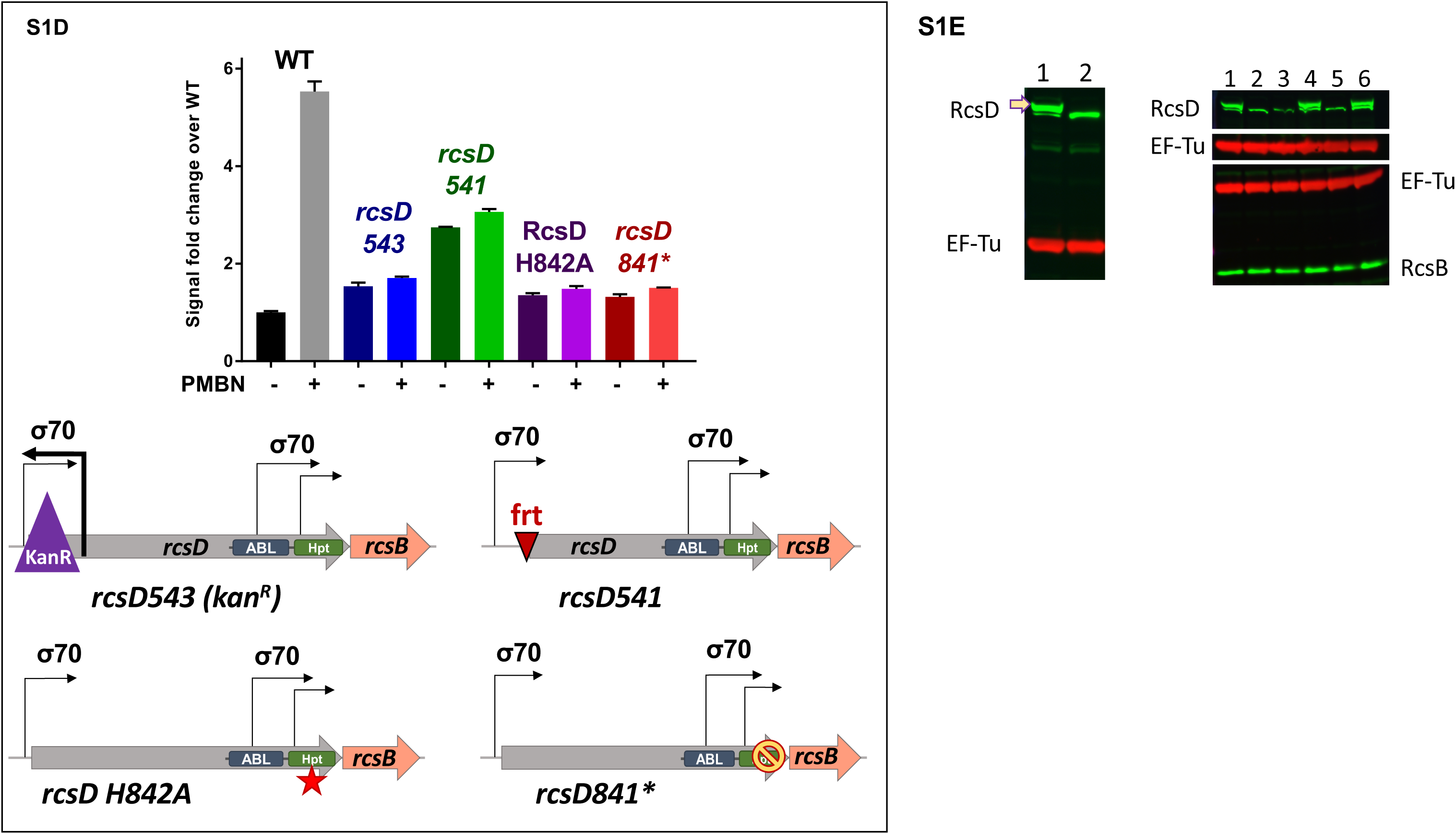
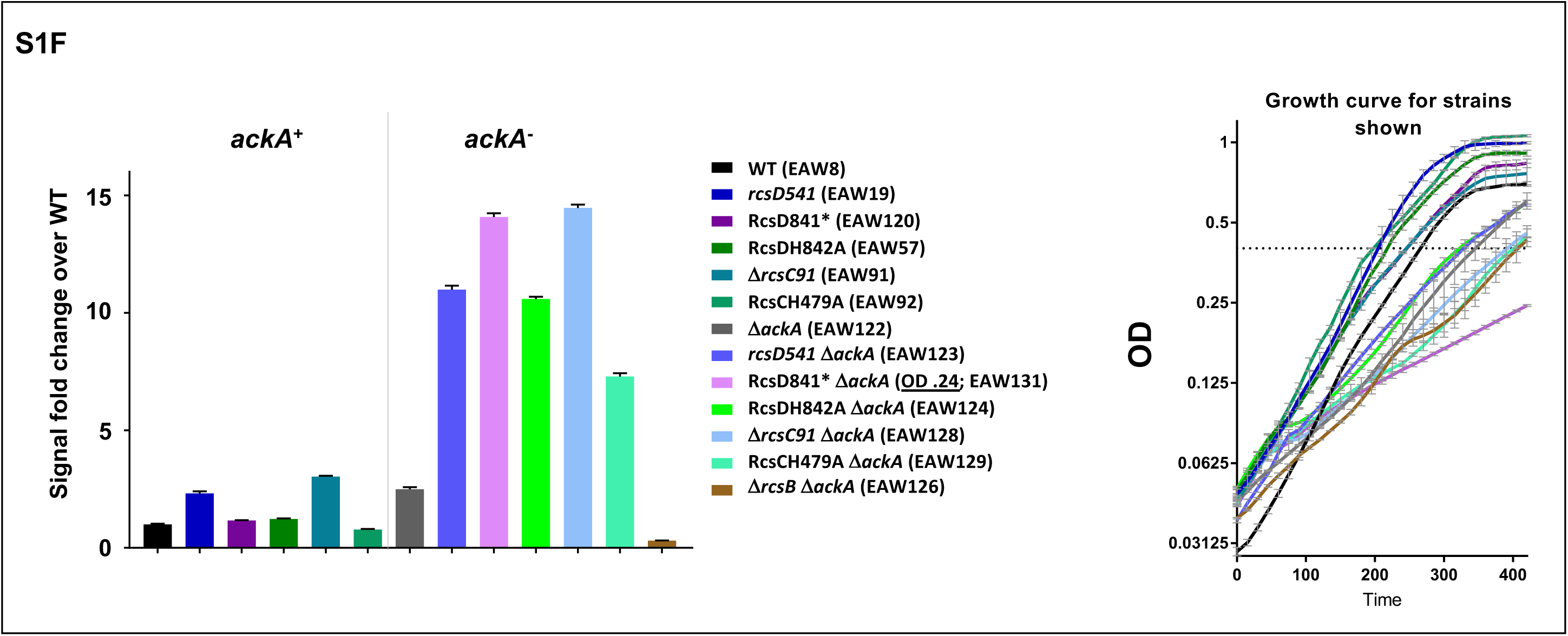
Measurement of Rcs activity by a fluorescent assay (relevant to Fig 1B) A) S1A-C use the same color code as Fig 1B, with strains and treatments listed in color code. Growth curve of each strain +/- PMBN 20 µg/mL as shown in Fig 1B. Dotted lines represent an OD_600_ of 0.4 and a 360 min (6 hour) time point, used as the standard measurements for fluorescent strains, unless stated otherwise. Demonstrated in A is that stationary phase doesn’t begin for any strain until close to or after OD_600_ 0.8 under plate reader growth conditions. Stationary phase always induces Rcs and can cause buildup of cells in well bottoms; therefore, measurements were not made past OD_600_ 0.8. If a strain has a growth defect that does not allow it to reach OD_600_ 0.4 before the 360 min time point, it is noted with its actual OD_600_ on the relevant bar graph legend in the figures. B) Relative fluorescent units (RFU) as a function of OD_600_ for strains used in Fig 1B. The vertical dotted line represents the measurement point that is shown in the Fig 1B bar graph, OD_600_ 0.4. These traces demonstrate the overall differences in Rcs activation of each strain. One can note the effect of PMBN +/- on the slope of each line. For example, WT without PMBN (black) has a low slope throughout the graph, while WT +PMBN (gray) has a noticeably higher slope. This can be contrasted with the *rcsC* or *rcsD* mutants (blue and green respectively), which may have slight differences in RFU between treated and untreated conditions at each growth point, but this doesn’t dramatically affect the overall slope of the trace, indicating that small fluorescence differences here do not represent activation of Rcs as a whole. In addition, when a strain stops growing (here referring to WT+PMBN gray line at OD_600_ near 0.8) and the fluorescence continues to increase, the slope of the line becomes much sharper, and we avoid using measurements in this range due to stationary phase Rcs activation of the *rprA* promoter, which must be distinguished from stimulus or drug-induced activation. C) This graph is a “zoomed in” version of S1B Fig with only WT, *rcsF*^-^ and *rcsB*^-^ strains, and demonstrates that the point of divergence between the treated and untreated lines can be another interesting proxy for detecting Rcs activation. True Rcs activation occurs in early growth points and is consistent over the growth of the strain (see WT). This is the case for an *rcsF* mutant (orange) which has a lower basal level of signal, but the PMBN treated condition demonstrates a consistently higher slope, with no trace overlap after OD_600_ of about 0.1. *rcsB* deletion (red) gives a low slope with no reaction to PMBN, showing almost complete trace overlap. D) Four different chromosomal *rcsD* mutants were examined and were found to have different effects on P_rprA_::mCherry activity. Strains shown here include WT (EAW8), *rcsD*543 (EAW9), *rcsD*541 (EAW19), *rcsD* H842A (active site Hpt domain mutant, EAW57) and *rcsD*841* (two stop codons replace codons 842 and 843; EAW120). These alleles are also depicted in the gene schematics. *rcsD*543 contains a non-polar Kanamycin resistance cassette that is transcribed in the opposite direction to *rcsD*; the Kan cassette deletes everything from the RBS to 540bp inside the *rcsD* ORF. Our most commonly used mutant, *rcsD*541, is a markerless deletion that results from Flp recombinase removal of the Kan cassette from a different construct, but it has the exact same deletion boundaries as *rcsD*543, with a frt scar and no reverse promoter. *rcsD*841* was expected to make a truncated protein, but in a Western blot is found to have no identifiable protein in the correct size range (S1E Fig). We have concluded that *rcsD*841* is probably a true RcsD null; the difference in expression with *rcsD*541 is intriguing but unexplained. *rcsD* H842A produces protein of the correct size, yet has the same level of P_rprA_-mCherry activation as deletion alleles 543 and 841*. None of these alleles appear to be polar on RcsB levels (S1E, right panel). E) Left blot: This polyclonal RcsD antibody can detect full length protein, but also detects a nonspecific band only slightly lower in molecular weight. 1) Wild type (EAW8), 2) complete deletion of RcsD ORF with the kan^R^ AraC Kid cassette (EAW52). Right blot: Parallel detection of RcsD and RcsB to check RcsD alleles for RcsB polarity. Underlined constructs produce RcsD at expected molecular weight. 1) <u>WT (EAW8)</u>, 2) *rcsD543* (EAW9), 3) *rcsD541* (EAW19), 4) <u>RcsD H842A (EAW57)</u>, 5) *rcsD841\** (EAW120), 6) <u>RcsD T411A (EAW121)</u>. F) An *ackA* mutant accumulates higher levels of acetyl phosphate, leading to phosphorylation of RcsB and thus activity of the P*_rprA_*-mCherry reporter. This increase is modest (two-fold) in a strain wild-type for the Rcs phosphorelay (WT; black and gray bars). The significantly higher activity in the *rcsC* and *rcsD* mutants is interpreted as a defect in dephosphorylation of RcsB∼P. Thus, *rcsD*541, 841* and H842A (blue (EAW123), purple (EAW131) and green (EAW124) bars) all lose the ability to dephosphorylate RcsB, easily evident in an *ack*A background. Although also unable to fully dephosphorylate RcsB∼P, a markerless whole-ORF *rcsC* deletion (EAW128; no RcsC receiver domain) and *rcsC* H479A (EAW129; intact RcsC receiver domain) appear to differ in their ability to perform the phosphatase reaction, consistent with existing literature about the primacy of receiver domains in the dephosphorylation reaction [16]. All *ackA* mutants have a slight growth defect (right panel); *rcsD*841* is the most defective. For this strain, the sample was taken at OD_600_ 0.24, possibly leading to an underestimate of its activity.

**S2 Fig.**
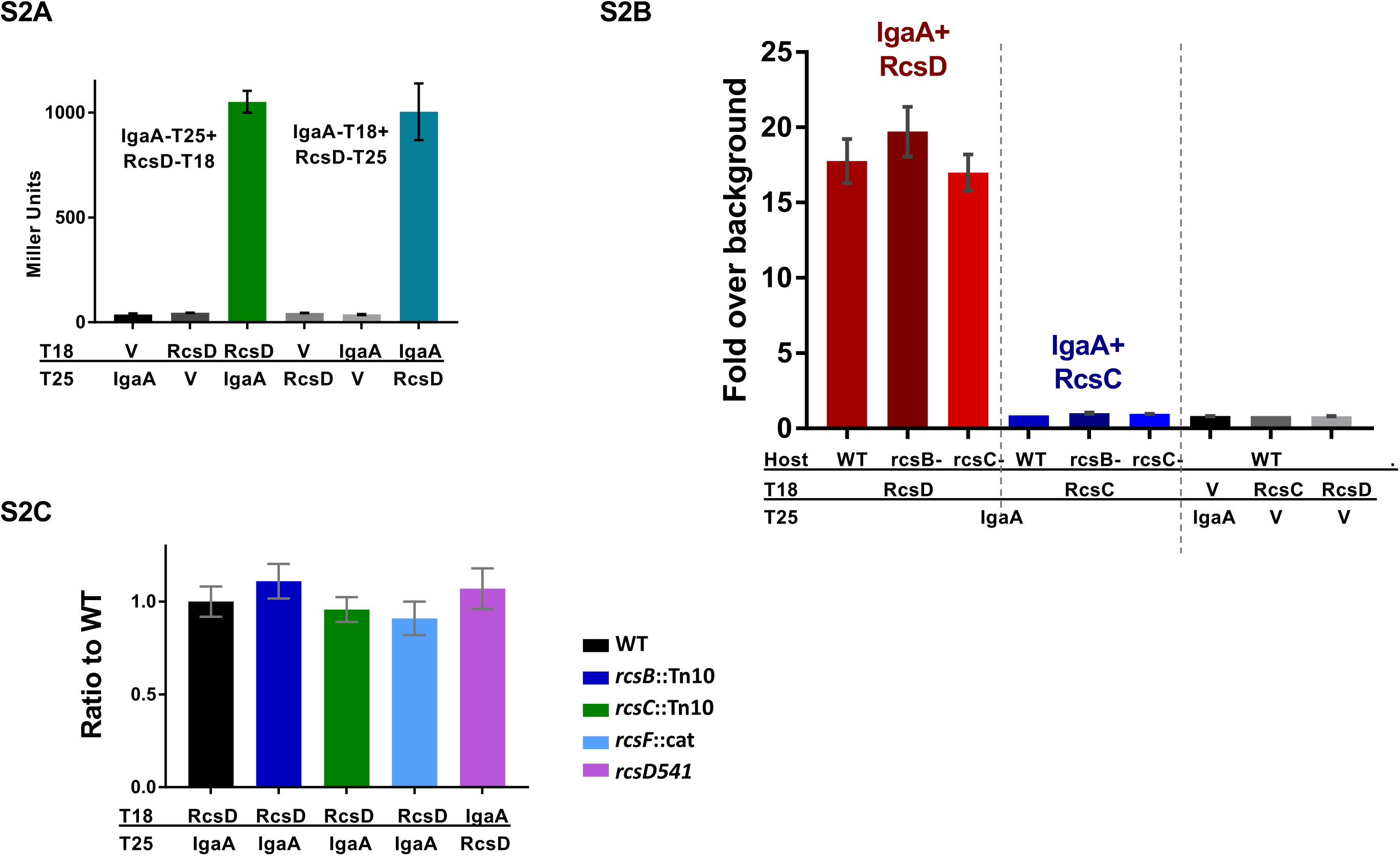
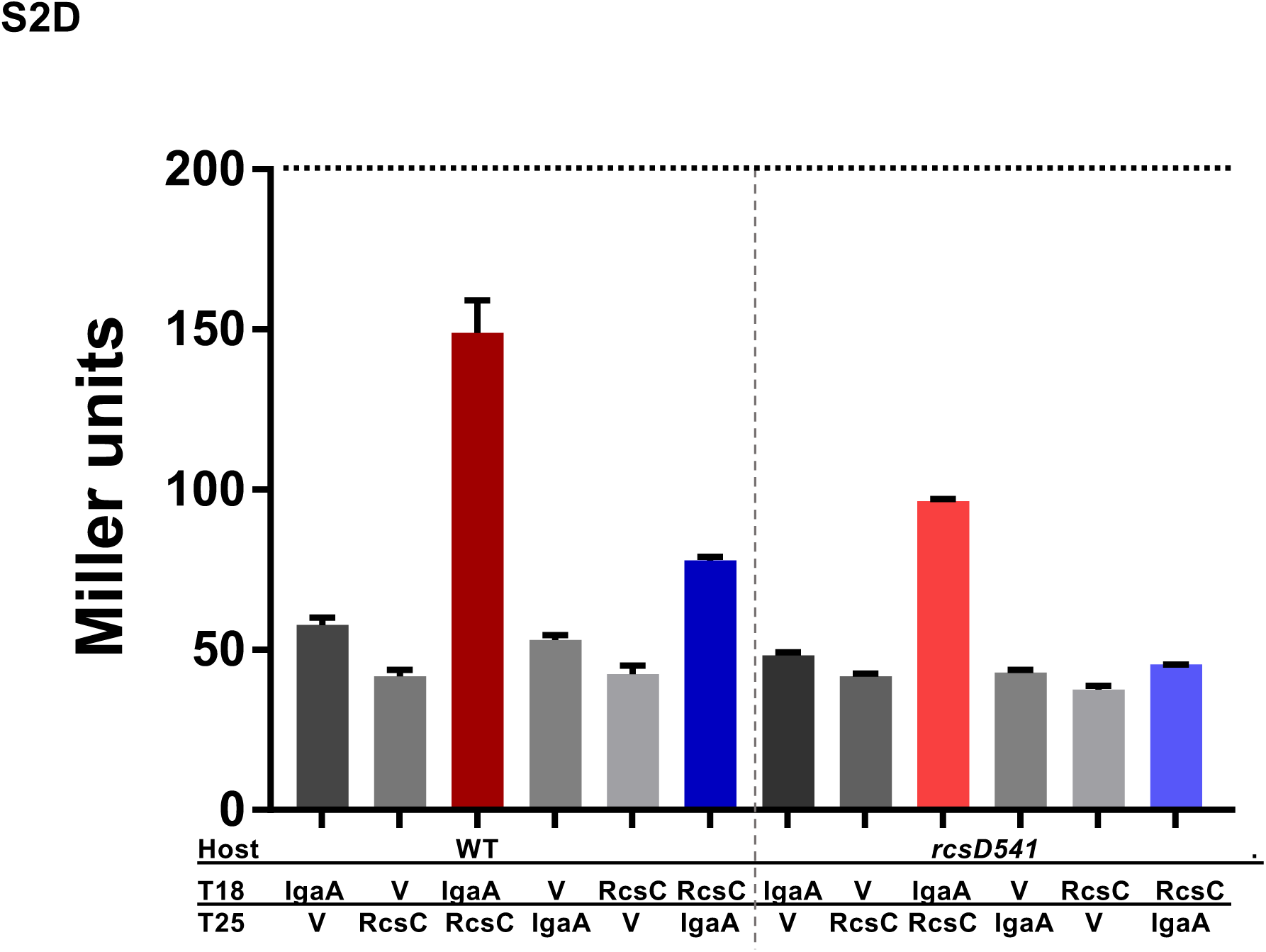
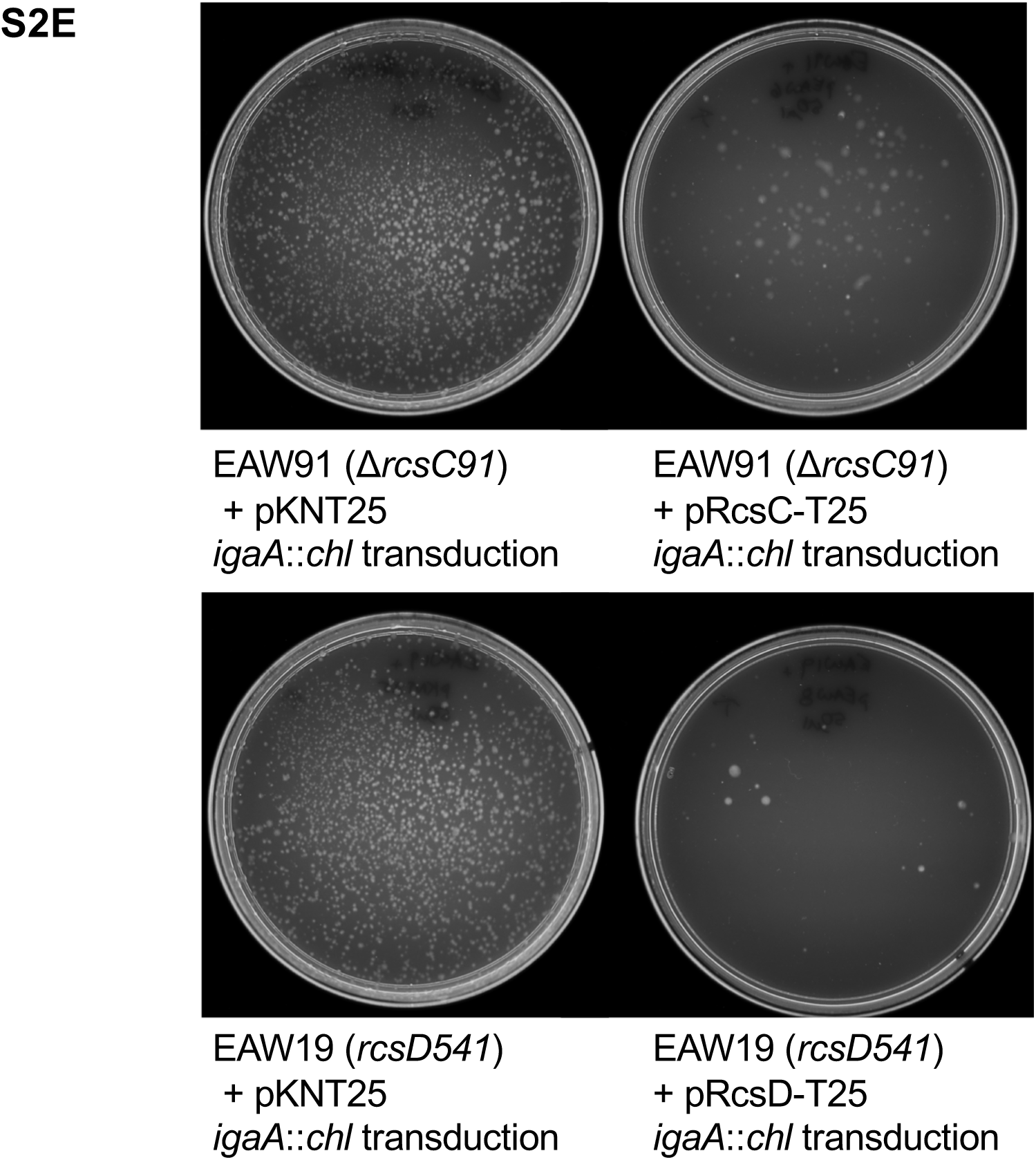
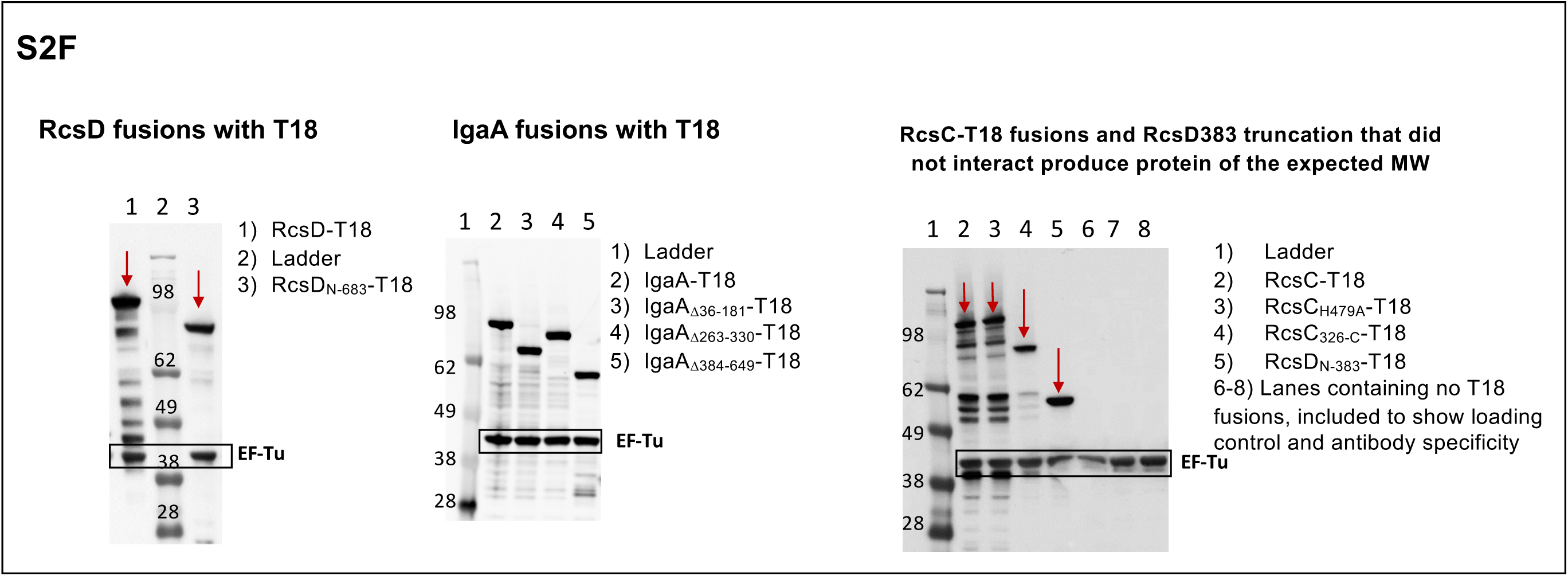
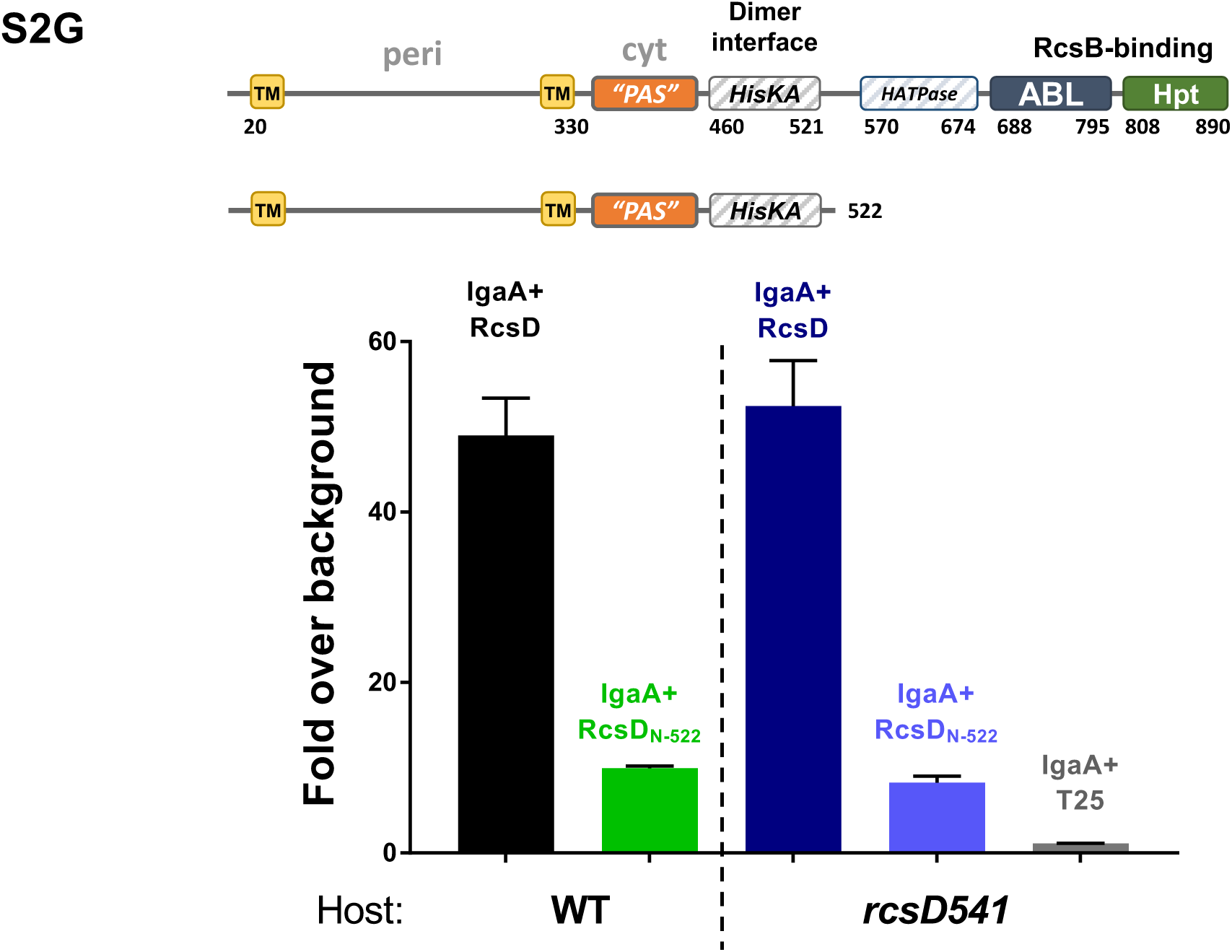
Interaction of IgaA and RcsD in a Bacterial Two-Hybrid Assay (relevant to Figure 2) A) IgaA and RcsD interact well regardless of which tag is used on each. The interaction registers at least 1000 Miller units, while vector control experiments yield only 50, giving a 20-fold signal to noise ratio, used in most graphs. Plasmids used: pEAW1 (IgaA-T18), pEAW2 (IgaA-T25), pEAW7 (RcsD-T25), pEAW8 (RcsD-T18). All error bars throughout the figures represent standard deviation. B) IgaA and RcsD interact approximately 20-fold over control empty vectors, regardless of strain background. IgaA/RcsC interaction is below the limit of detection in all strains tested. Empty vector controls were performed in the WT background (BTH101), *rcsB*::Tn10 (EAW1), and *rcsC*::Tn10 (EAW2). C) Results from S2B merged with results from different experiments done in the *rcsF*^-^ (EAW4) and *rcsD*^-^ (EAW12) backgrounds. Each bar represents the relative IgaA/RcsD interaction measurement in the respective mutant host relative to the IgaA/RcsD interaction in wild type cells; this positive control is present for normalization in every assay of RcsD or IgaA variant interaction. D) RcsC interaction with IgaA cannot be reliably detected irrespective of tag orientation or strain background. IgaA/RcsC were fused in both orientations and tested in WT (BTH101) and *rcsD541* (EAW12) host backgrounds. The dotted line at 200 Miller units represents approximately 4-fold over the background controls, the standard for a consistent, repeatable interaction determination. Note difference in beta-galactosidase values for even the strongest interaction here (150 Miller units) compared to the interaction of RcsD with IgaA (S2A Fig). Plasmids used: pEAW1 (IgaA-T18), pEAW6 (RcsC-T25), pEAW2 (IgaA-T25), and pEAW5 (RcsC-T18). V: vector, pUT18 for the T18 vector and pKNT25 for T25 vector. E) RcsD-T25 and RcsC-T25 fusions are functional, as judged by complementation of chromosomal mutations for lethality in the absence of IgaA. When the *rcsC* strain EAW91 and *rcsD*541 strain EAW19 containing empty vector are transduced with a chloramphenicol resistant *igaA* deletion allele from NM357 (*igaA*::Chl), many colonies result (left plate in each pair), because IgaA is only essential when the Rcs system is able to actively signal. When these strains contain RcsC-T25 (pEAW6) or RcsD-T25 (pEAW8) respectively, the Rcs signaling cascade is restored and deletion of *igaA* is no longer possible (right plate in each pair), demonstrating functionality of the RcsC-T25 and RcsD-T25 constructs. Rare colonies that do result on these plates are mucoid and/or mutant. F) Western blot of T18 fusion proteins, using the T18 antibody in a *cya*+ strain (NEB Turbo), regardless of whether the proteins form detectable interactions. Red arrows show the expected fusion proteins in blots with multiple bands. Plasmids present are pEAW7 (RcsD-T18), pEAW7b (RcsD_N-683_-T18), pEAW1 (IgaA-T18), pEAW1cyt1 (IgaA_Δ36-181_-T18), pEAW1cyt2 (IgaA _Δ263-330_-T18), pEAW1peri (IgaA _Δ384-649_-T18), pEAW5 (RcsC-T18), pEAW5H (RcsC_H479A_-T18), pEAW5s (RcsC_326-C_-T18), pEAW7m (RcsD_N-383_-T18). G) RcsD522 is not deficient in binding IgaA due to interaction with chromosomal RcsD. Here the interaction of FL RcsD with IgaA and the interaction of RcsD_N-522_ with IgaA was unaffected by mutation of rcsD (compare left, WT, to right, rcsD541 strain background). NB: the signal for IgaA/RcsD interaction was 50-fold above background here, as opposed to 20-fold (only one background control is shown for brevity; as in previous figures, background controls do not vary significantly from one another). Present are pEAW1 (IgaA-T18), pEAW8 (RcsD-T25), and pEAW8alpha (RcsD_N-522_-T25) in hosts BTH101 (WT) and EAW12 (*rcsD451*).

**S3 Figure:**
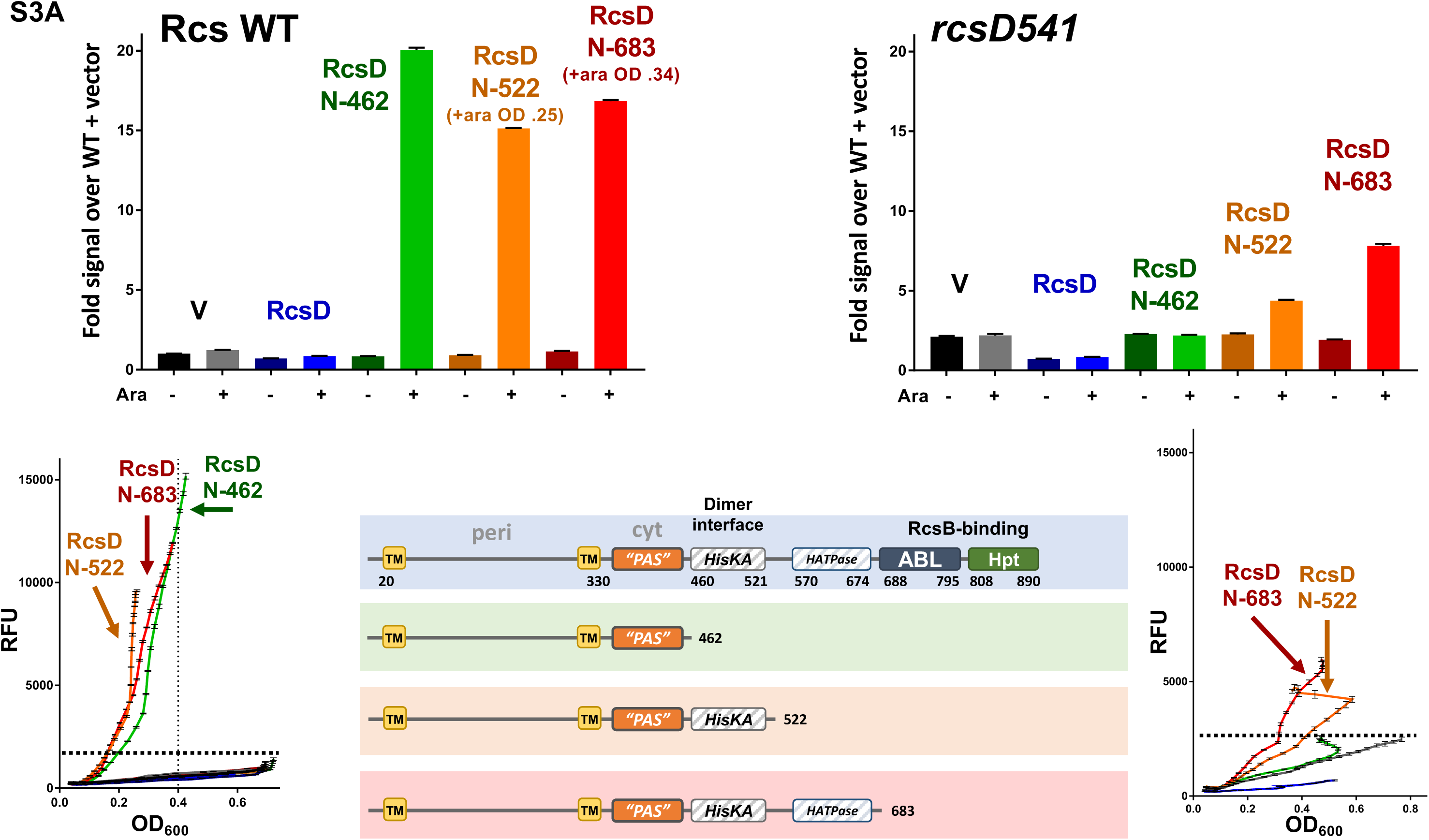
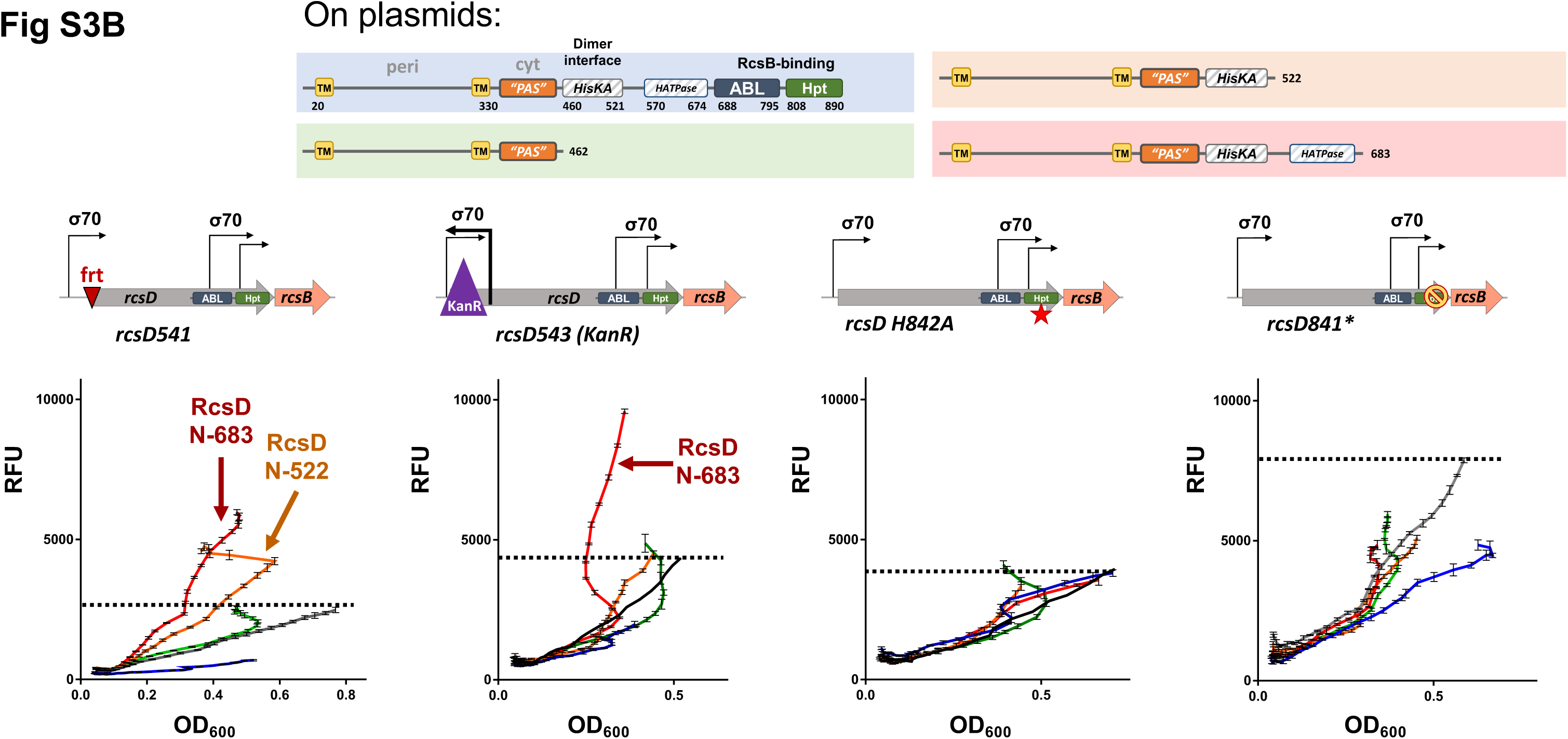
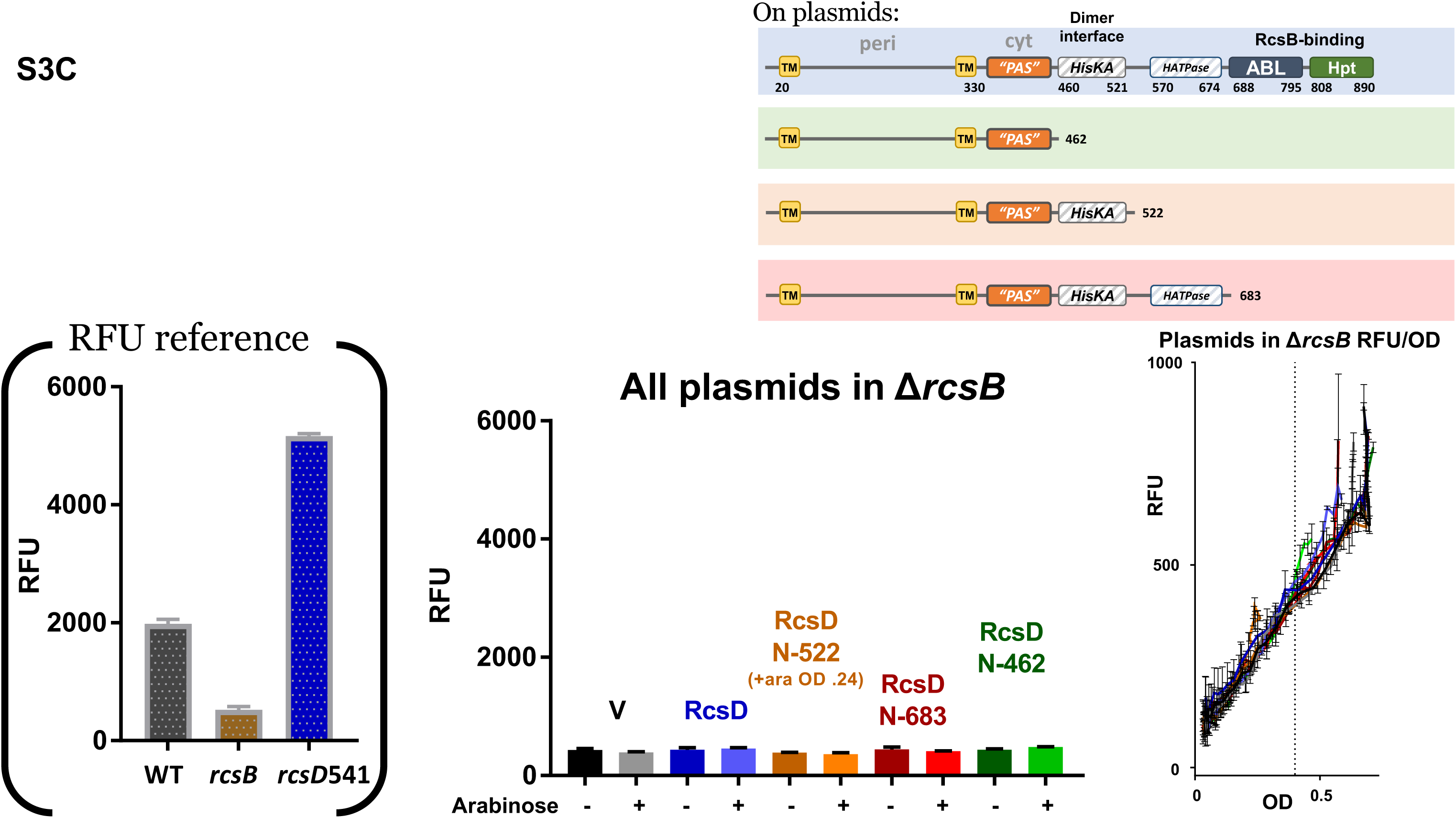
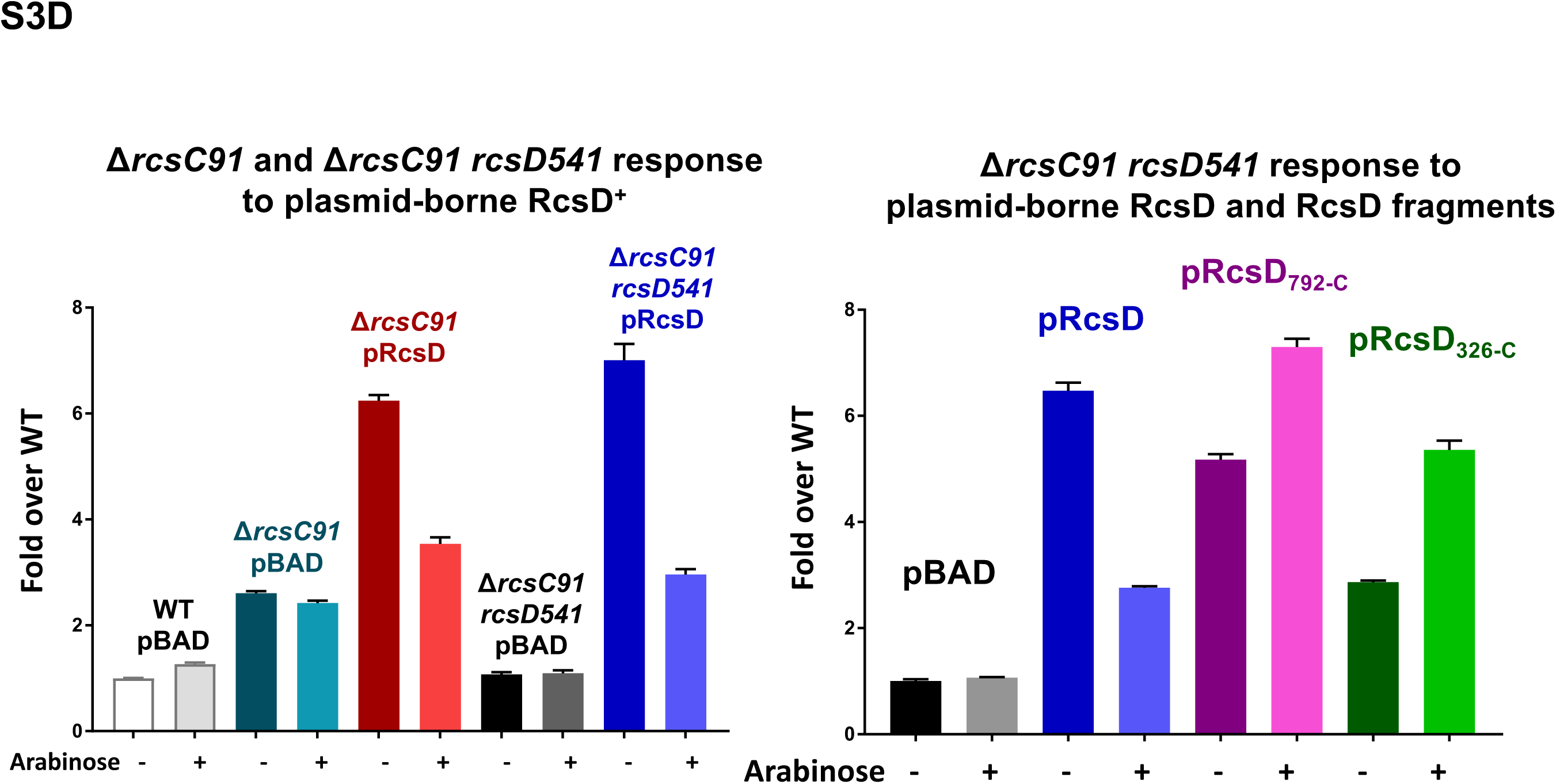
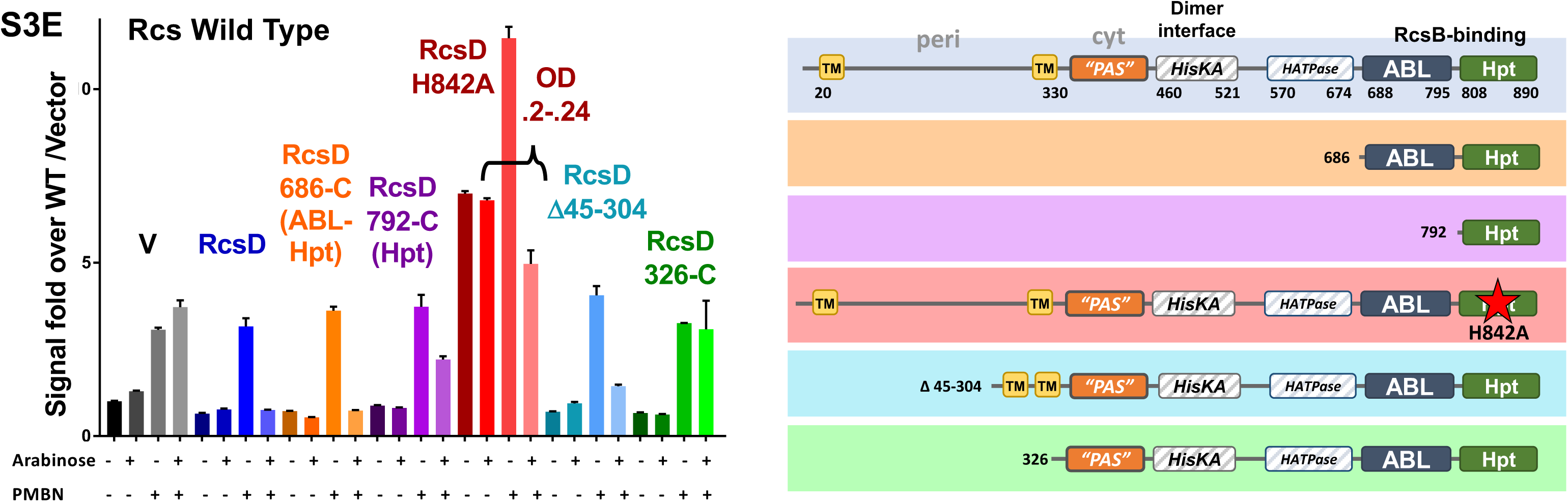
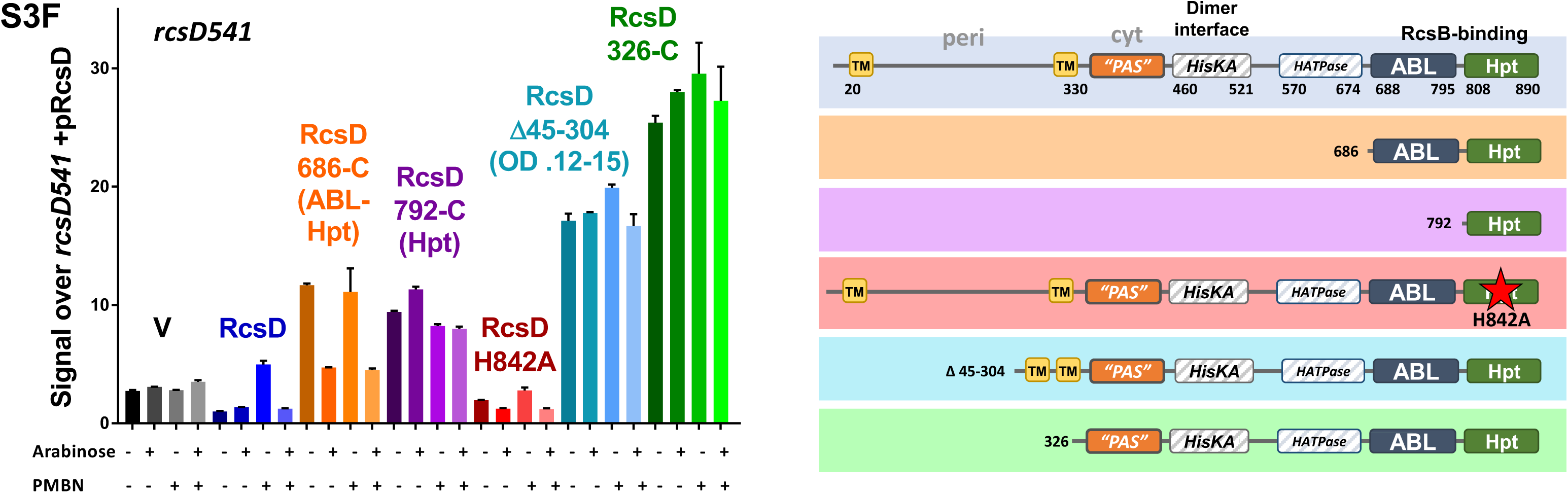
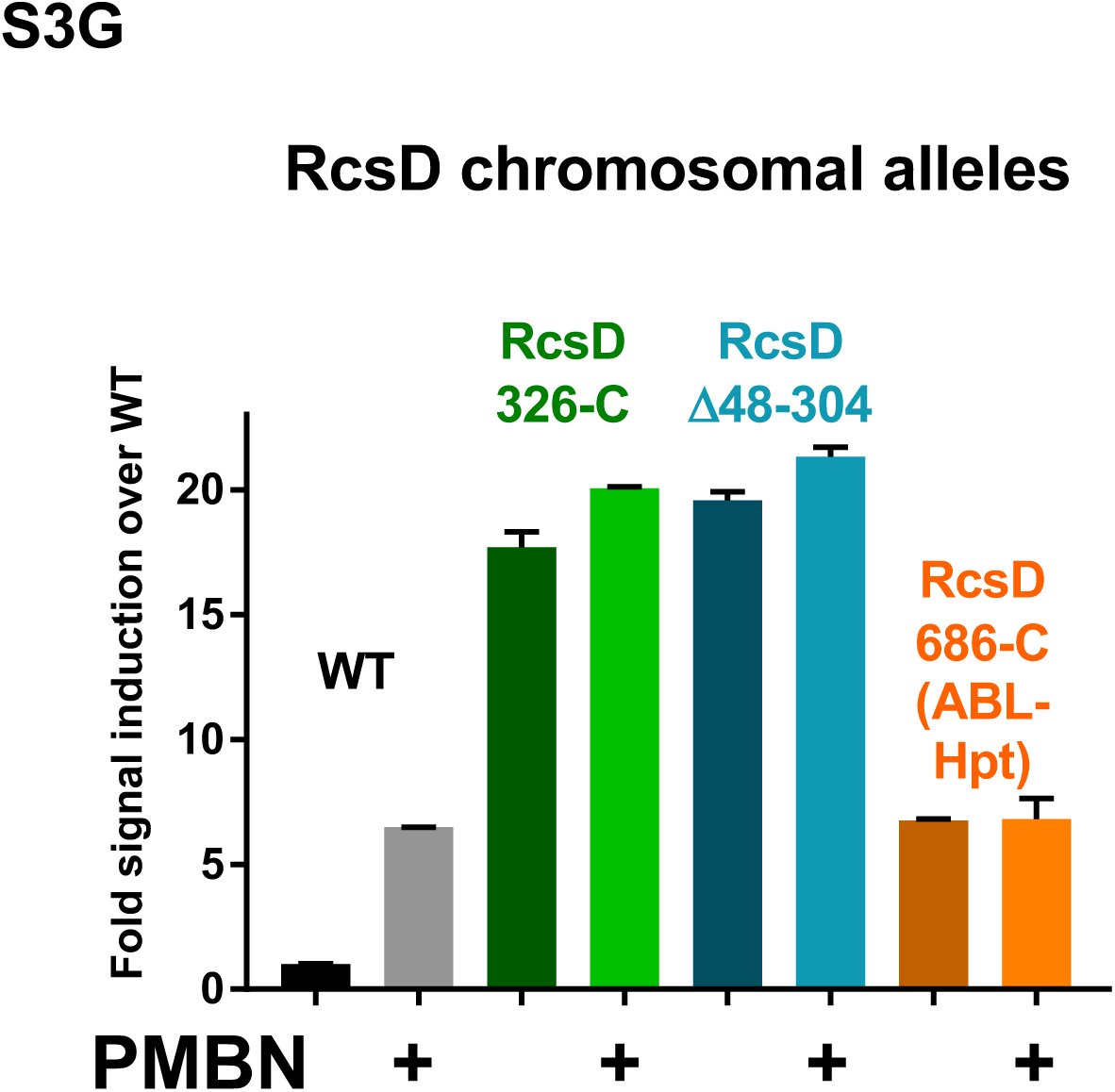
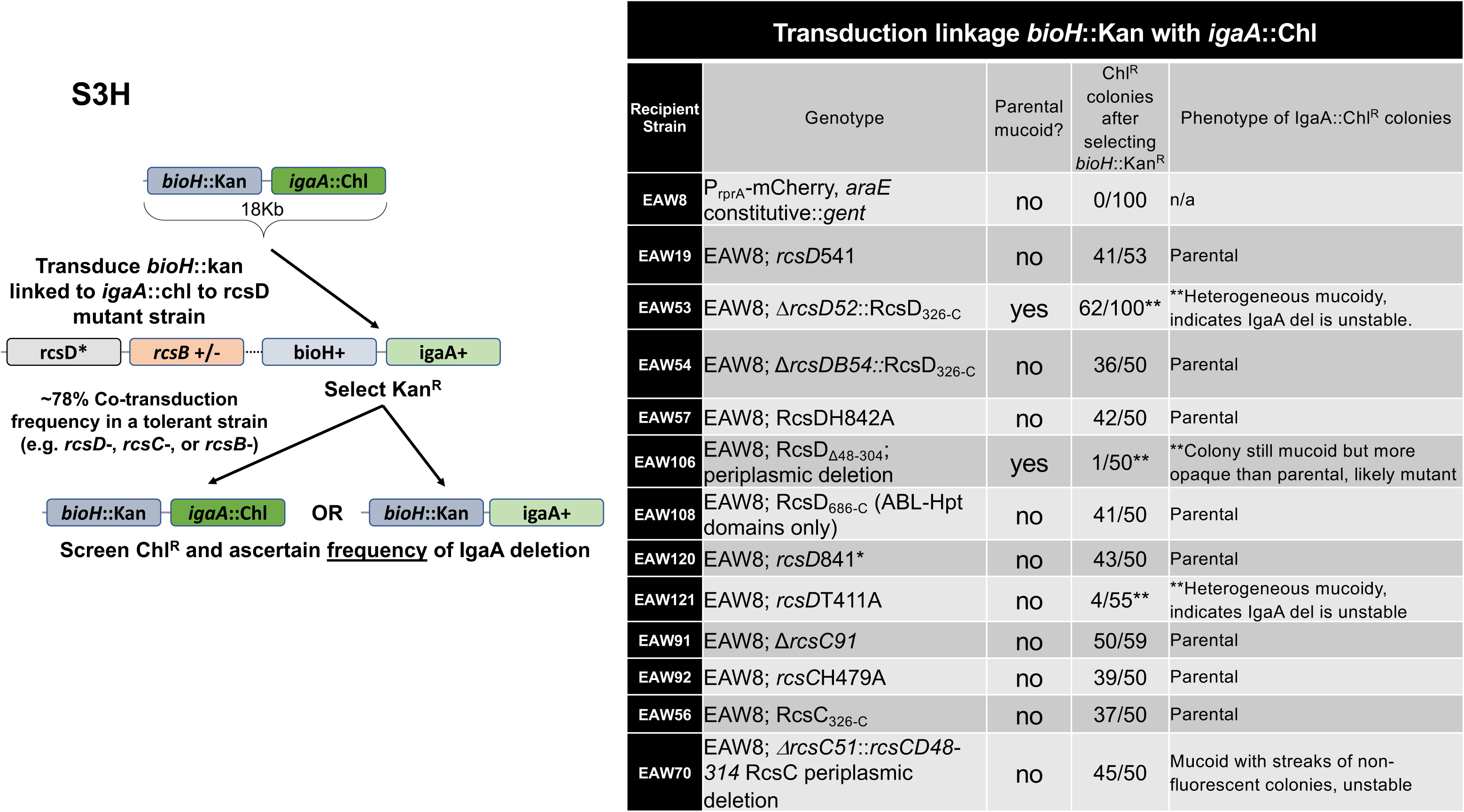
Analysis of RcsD function in signaling. **A.** As for Fig 3A, with additional plasmids. RcsD C-terminal truncation constructs were expressed from arabinose-inducible plasmids in a WT (EAW8) and a *rcsD*541 (EAW19) host. The graphs of strain fluorescence as a function of OD_600_ for cells grown with arabinose are also presented below their respective bar graphs. Constructs are color-coded: black: V, (pBAD24); blue: RcsD^+^, (pEAW11); green: RcsD_N-462_ (pEAW11m2); orange: RcsD_N-522_ (pEAW11*α*); red: RcsD_N-683_ (pEAW11b). Note that a change in slope on the fluorescence/ OD_600_ graph demonstrates some level of P*_rprA_*-mCherry activation, and that the orange (RcsD_N-522_) and red (RcsD_N-683_) slopes are very different than other slopes in the *rcsD*541 strain. Cell lysis can be seen as a reduction in OD_600_ resulting in a leftward shift in the line (see orange and green lines in *rcsD*541 host). Note that, in spite of lysis for RcsD_N-462_ in *rcsD*541, greater fluorescence did not result, compared to the vector control in the same time period. Therefore, lysis does not automatically increase P*_rprA_*-mCherry fluorescence. Highest RFU with vector shown by horizontal dotted line, for comparison with experimental curves. **B.** Activity of plasmids in different *rcsD* mutants. Based on the unexpected signal from plasmids lacking the Hpt domain in *rcsD*541 (S3A Fig), further *rcsD* alleles were tested with RcsD C-terminal truncation plasmids. Fluorescence as a function of OD_600_ is shown for cells grown as in S3A Fig, but in strains carrying four different chromosomal *rcsD* alleles, *rcsD*541 (EAW19, repeated from S3A Fig), *rcsD*543 (EAW9), *rcsD*H842A (EAW57) and *rcsD*841* (two stop codons at residue 841, EAW120). Each *rcsD* allele is shown as an inset above the Fluorescence/ OD_600_ trace for that strain. Plasmids are color-coded as in S3A Fig. RcsD_N-522_ and RcsD_N-683_ in the *rcsD*541 and *rcsD*543 strains achieve higher slopes and/or final RFU values than the vector control. Highest RFU with vector shown by horizontal dotted line, for comparison with experimental curves. The same is not true in backgrounds containing a disrupted Hpt domain. How activation is occurring in *rcsD*541 and *rcsD*543 is unexplained, but it apparently requires the presence of the RcsD Hpt domain in the chromosome. **C.** Overexpression of RcsD C-terminal truncations cannot activate in the absence of RcsB, demonstrating that their effect on P*_rprA_*-mCherry is Rcs pathway specific. Assays and color-coding are as in S3A Fig, but in an *rcsB*::kan strain (EAW31). Shown here (L to R) are a bar graph with *rcsB* RFU compared to WT, a bar graph (OD_600_ 0.4 or final OD_600_ value at 6 hours) where the RFU values for each construct can be easily visualized and compared, and a graph of relative fluorescence units as a function of OD_600_. There are no significant differences in slope or final RFU value, and the RFU values are the same as the background levels of P*_rprA_*-mCherry expression in an *rcsB* deletion. **D.** Overexpression of RcsD constructs containing the Hpt domain depend on RcsC for highly unregulated activation and mucoidy. In the left bar graph, RcsD on a plasmid is compared to empty vector in WT, Δr*csC* (EAW91) and Δr*csC rcsD*541 (EAW93) strains. Although signal increases in both *rcsC* deletion backgrounds, there is no mucoidy in these strains. The threshold for mucoidy is closer to twelve-fold over wildtype; these strains approach seven-fold. In the right bar graph color-coding is as in Fig 3B, but all plasmids are in strain EAW93. RcsC is necessary for mucoidy, but apparently not for smaller signal increases. **E, F.** Strains (E.: EAW8, WT; F: EAW19, *rcsD*541) containing the indicated RcsD plasmids were grown and assayed under four conditions: without arabinose or PMBN, with arabinose (.02%), with PMBN, and with arabinose and PMBN. **G.** *rcsD* alleles were introduced into the chromosomal *rcsD* locus to create: *rcsD*_326-C_ (EAW53), *rcsD*_Δ48-304_ (EAW106) and *rcsD*_686-C_ (EAW108). *rcsD*_792-C_ could not be introduced without deleting promoters for RcsB, so that construct was not made. These alleles performed as their plasmid counterparts did, with the longer constructs roughly equivalent in their high signal and slow growth and the *rcsD*_686-C_ allele appearing less efficient at passing signal to RcsB. None of these alleles respond to PMBN, but only the *rcsD*_686-C_ allele can tolerate an *igaA* deletion (S3H Fig). **H.** Co-transduction of *igaA*::chl^R^ with *bioH*::kan as an assay of Rcs function. Schematic shows *igaA*::chl^R^ cotransduction frequency experiment using linked *bioH*::kan. The *bioH*::kan *igaA*::chl^R^ donor (EAW66) was constructed in an *rcsD* mutant. Table lists frequency of *igaA*::chl^R^ cotransduction into various *rcs* mutants, all isogenic derivatives of the Rcs^+^ strain EAW8, noting the phenotype of transductants.

**Fig. S4:**
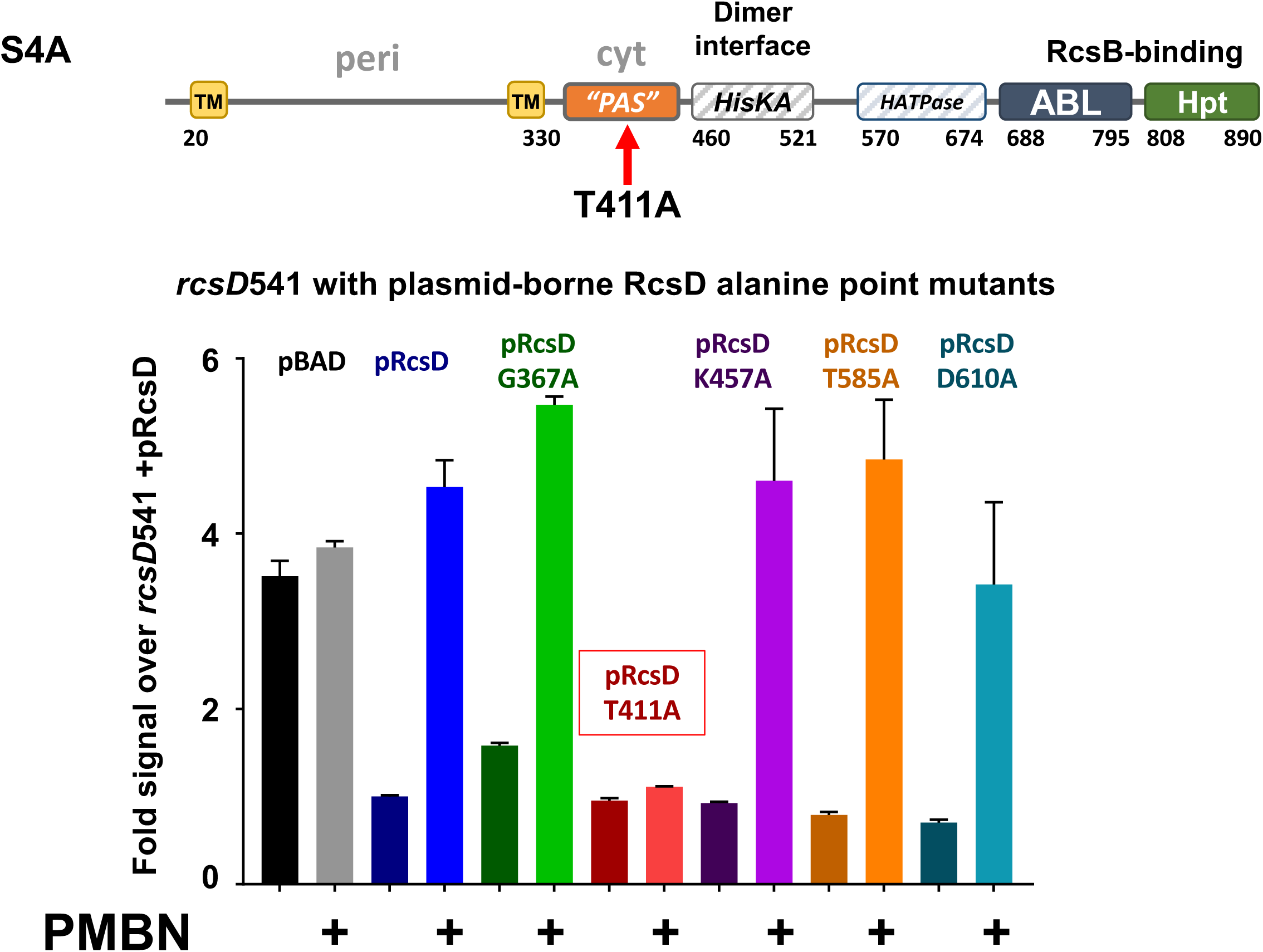
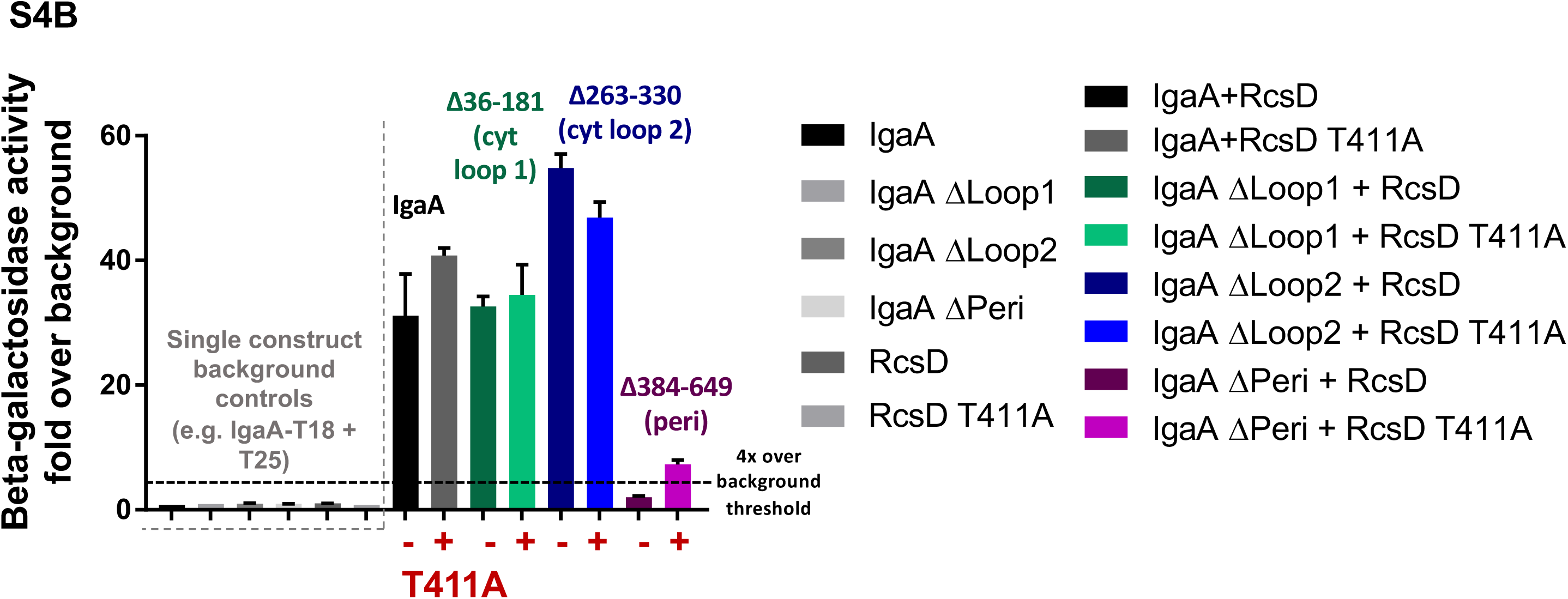
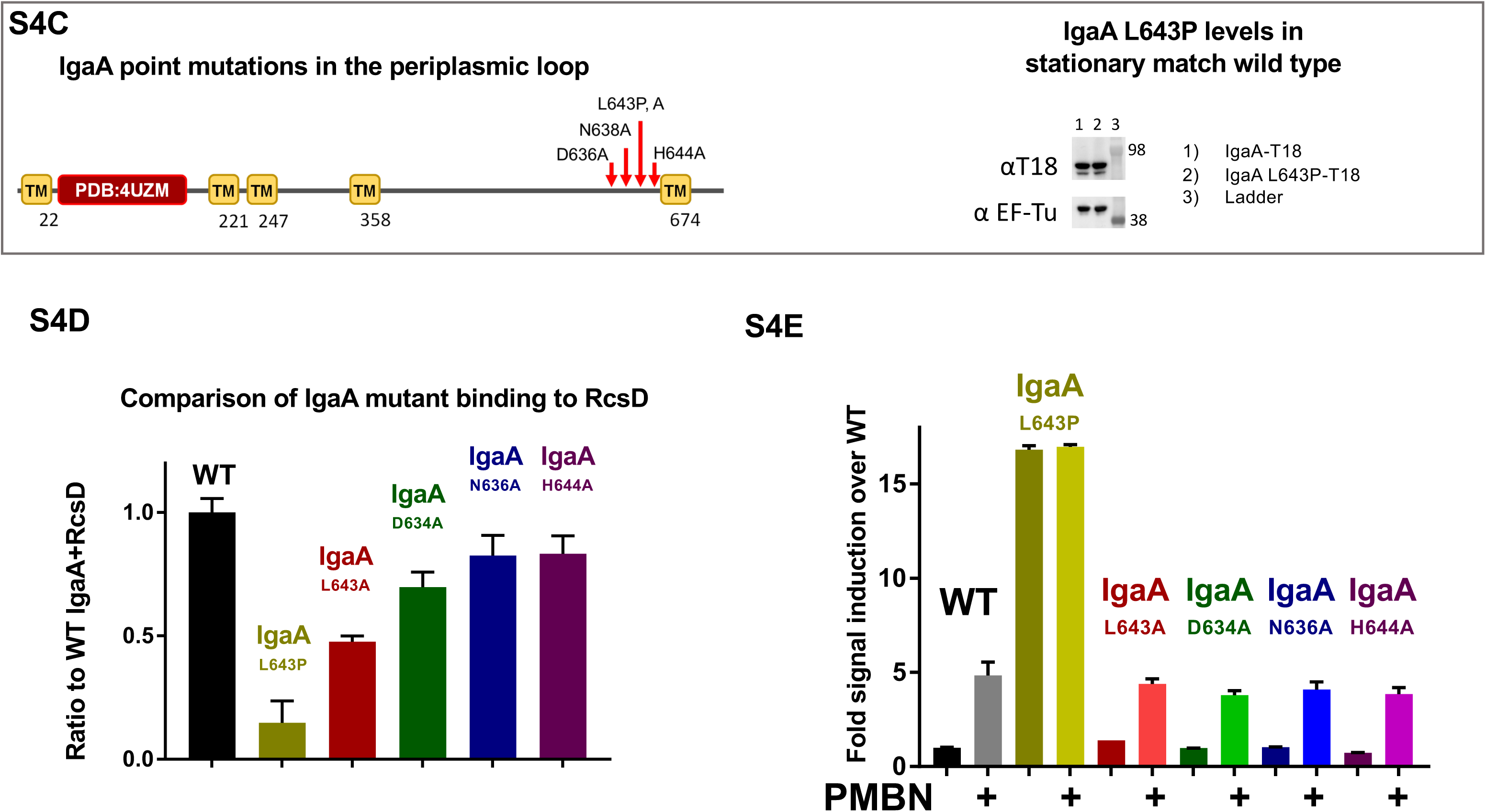
Mutation in RcsD cytoplasmic region blocks signaling. A. Strain EAW19 (*rcsD*541) with derivatives of pBAD24-RcsD (pEAW11) grown in MOPS glucose or MOPS glucose with PMBN, grown without arabinose and assayed as in S3E, F Fig. *rcsD541* has a higher signal than wild type, and can be complemented with WT RcsD on a plasmid (compare lanes 1 and 3). The RcsD^+^ construct responds to PMBN, unlike empty vector (compare lanes 1 to 2 and 3 to 4). Plasmids encoding *rcsD* alanine mutations in the cytoplasmic domains were screened for those that complemented *rcsD*541, reducing the basal level of expression; these were then assayed with and without PMBN. Of the 5 alleles shown here, 4 were inducible with PMBN. However, although it complements an *rcsD* deletion by lowering signal, the point mutant *rcsD*T411A did not respond to PMBN. B. BACTH IgaA loop deletion interactions with RcsD WT vs RcsD T411A, as in Fig 4B, but showing the various controls. The IgaA+RcsD constructs give signal greater than thirty-fold over the single construct (background) controls. T18 derivatives carrying IgaA cytoplasmic loop one deletion (Δ36-181, cyt loop 1; pEAW1cyt1), IgaA cytoplasmic loop two deletion (Δ 263-330, cyt loop 2; pEAW1cyt2), IgaA periplasmic loop deletion (Δ 384-649, peri; pEAW1peri) were tested with RcsD-T25 WT (pEAW8) or RcsD T411A (pEAW8T) were tested in BTH101. C. Schematic showing point mutations surrounding IgaA L643P, a mutant of IgaA defective in Rcs negative regulation. In a western blot, the level of the T18-IgaA fusion protein is similar for L643P and wild type IgaA, indicating stability. EF-Tu was used as a loading control. Plasmids present: pEAW1, pEAW1L. D. Wild type IgaA interaction with RcsD in the BACTH system is set to one, and compared to IgaA point mutant interactions with RcsD. IgaA L643P is deficient, but L643A is significantly better and surrounding mutants are nearly WT for RcsD interaction. Plasmids tested (L to R): pEAW1, pEAW8, pEAW1L, pEAW1LA, pEAW1D, pEAW1N, pEAW1H. E. When inserted into the chromosome in place of the wild-type *igaA* gene, only IgaA L643P produces Rcs dysregulation; the other mutants are wild-type for Rcs negative regulation and response to PMBN. Strains present (L to R): EAW8, EAW111, EAW112, EAW109, EAW110, EAW113.

**S5 Figure:**
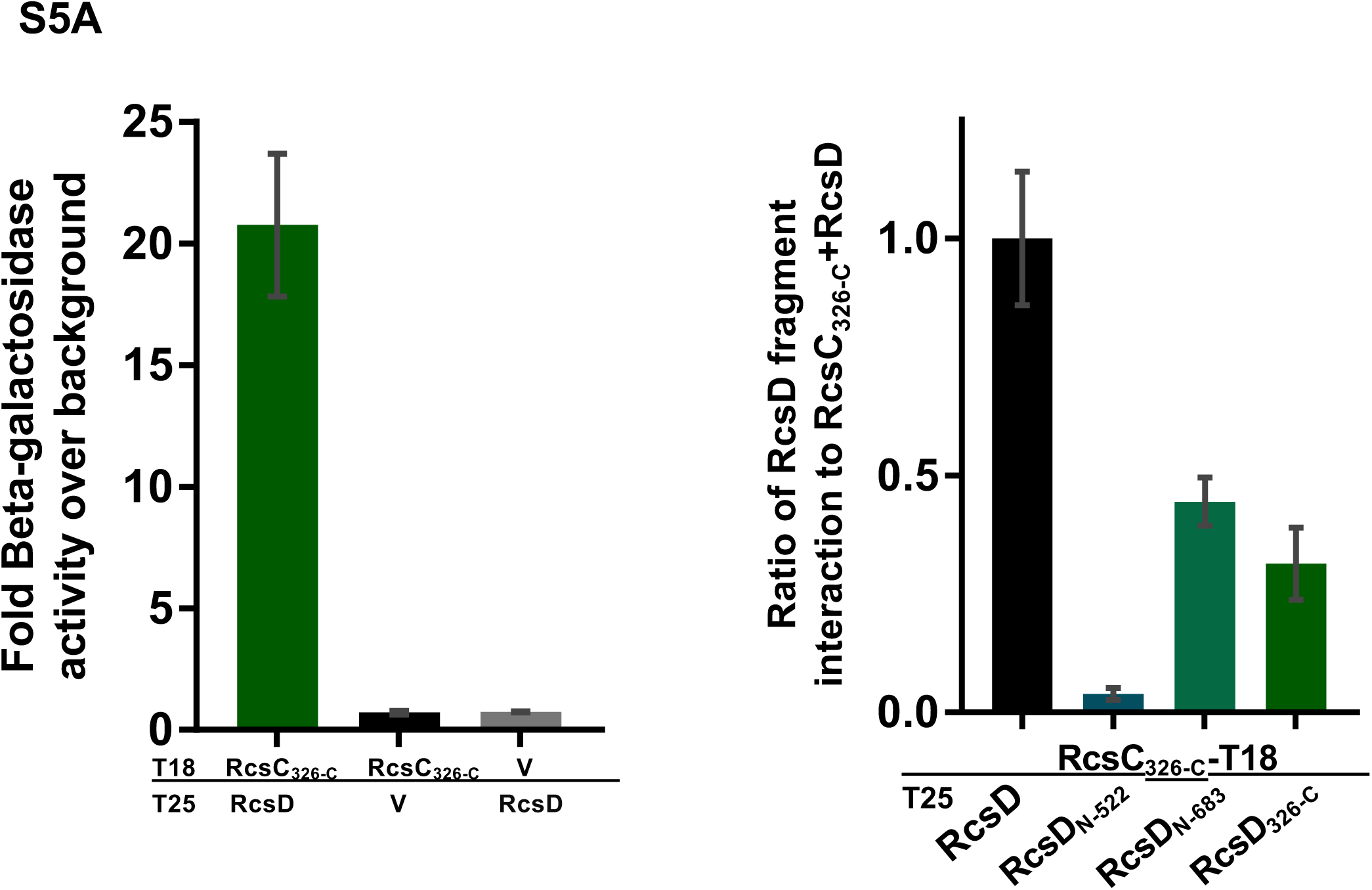
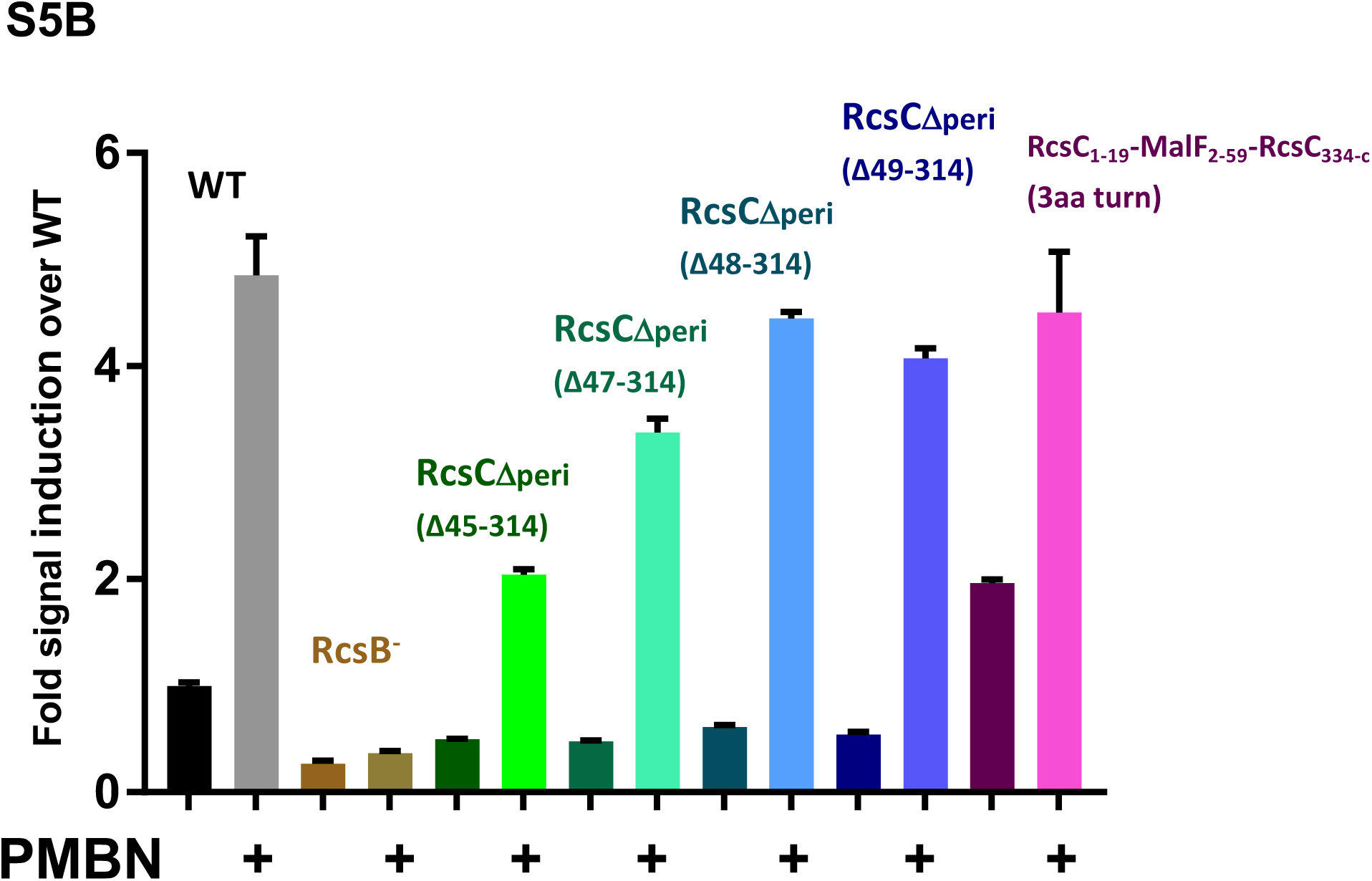
The RcsC periplasmic loop is dispensable for signaling. S5A: Bacterial two-hybrid assay of interaction of cytoplasmic portion of RcsC with regions of RcsD. Plasmids present include (L to R) pEAW5s, pEAW8, pEAW8α, pEAW8b, pEAW8s. S5B: RcsC periplasmic deletions perform differently when exposed to PMBN depending on the linker length between transmembrane domains and the identity of those transmembrane domains. Strains present include (L to R) EAW8, EAW31, EAW61, EAW69, EAW70, EAW71, EAW72.

**S6 Figure:**
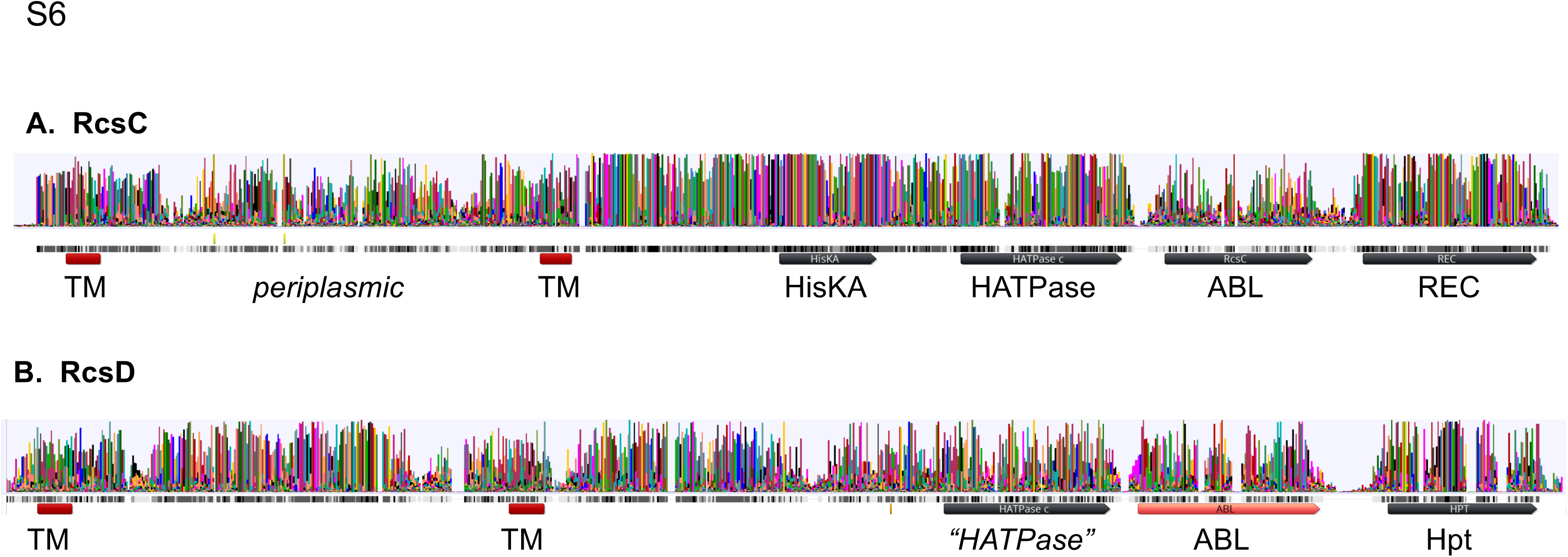
Conservation in RcsC and RcsD. A. Amino acid logo diagram for RcsC, created by aligning 77 RcsC proteins from different bacterial species within Enterobacterales. Conservation is high in the enzymatic regions (HisKA, HATPase, and REC domains), but is relatively low in the periplasmic region, found here to be dispensable for function and signaling. B. Amino acid logo for RcsD, created by aligning 83 RcsD proteins from different bacterial species within Enterobacterales. Conservation is not high in the ancestral histidine kinas and ATPase regions, but is high in the periplasmic domain. Alignments and logos were created in Geneious using MUSCLE.

### Supplemental Tables

S1 Table: Strains used in this study.

S2 Table: Plasmids used in this study.

S3 Table: Primers used in this study.

